# Illuminating a Phytochrome Paradigm – a Light-Activated Phosphatase in Two-Component Signaling Uncovered

**DOI:** 10.1101/2020.06.26.173310

**Authors:** Elina Multamäki, Rahul Nanekar, Dmitry Morozov, Topias Lievonen, David Golonka, Weixiao Yuan Wahlgren, Brigitte Stucki-Buchli, Jari Rossi, Vesa P. Hytönen, Sebastian Westenhoff, Janne A. Ihalainen, Andreas Möglich, Heikki Takala

**Affiliations:** Faculty of Medicine, Anatomy, University of Helsinki, 00014 Helsinki, Finland; Nanoscience Center, Department of Biological and Environmental Sciences, University of Jyvaskyla, 40014 Jyvaskyla, Finland; Nanoscience Center, Department of Chemistry, University of Jyvaskyla, 40014 Jyvaskyla, Finland; Lehrstuhl für Biochemie, Universität Bayreuth, 95447 Bayreuth, Germany; University of Gothenburg, Department of Chemistry and Molecular Biology, 40530 Gothenburg, Sweden; Faculty of Medicine and Health Technology, BioMediTech, Tampere University, 33520 Tampere, Finland; Fimlab Laboratories, 33520 Tampere, Finland

**Keywords:** phosphatase, phytochrome, response regulator, sensory photoreceptor, structural biology, two-component signaling

## Abstract

Bacterial phytochrome photoreceptors usually belong to two-component signaling systems which transmit environmental stimuli to a response regulator through a histidine kinase domain. Phytochromes switch between red light-absorbing and far-red light-absorbing states. Despite exhibiting extensive structural responses during this transition, the model bacteriophytochrome from *Deinococcus radiodurans* (DrBphP) lacks detectable kinase activity. Here, we resolve this long-standing conundrum by comparatively analyzing the interactions and output activities of DrBphP and a bacteriophytochrome from *Agrobacterium fabrum* (AgP1). Whereas AgP1 acts as a conventional histidine kinase, we identify DrBphP as a light-sensitive phosphatase. While AgP1 binds its cognate response regulator only transiently, DrBphP does so strongly, which is rationalized at the structural level. Our data pinpoint two key residues affecting the balance between kinase and phosphatase activities, which immediately bears on photoreception and two-component signaling. The opposing output activities in two highly similar bacteriophytochromes inform the use of light-controllable histidine kinases and phosphatases for optogenetics.

## INTRODUCTION

Two-component signaling systems are mainly found in prokaryotes and allow cells to respond to environmental signals^1^. These systems have been under extensive research ever since their discovery, as they control a wide range of cellular mechanisms from enzymatic activity to transcription regulation^2^. A canonical two-component system consists of a homodimeric sensor histidine kinase (HK) and its cognate response regulator (RR)^3^. To the extent it has been studied, most HK proteins sense chemical signals and generally reside within the plasma membrane^2^. The output activity is exerted by an intracellular HK module, consisting of two subdomains: a dimerization histidine phosphotransfer (DHp) domain, and a catalytic ATP-binding (CA) domain. Based on their DHp sequence, the HK proteins can be divided into four subtypes, called HisKA, HisKA_2, HWE_HK and HisKA_3^4,5^.

The HK catalyzes autophosphorylation and subsequent phosphotransfer to the cognate RR. During the autophosphorylation reaction, the eponymous histidine of the DHp domain is phosphorylated^6,7^, either within the same monomer (*cis*) or the sister molecule of the homodimer (*trans*)^8^. In the phosphotransfer reaction, the phosphate is relayed to a conserved aspartate residue within a receiver (REC) domain of the RR. This reaction entails RR activation and elicits output responses such as altered gene expression^3,6,9^.

HKs may also act as phosphatases that hydrolyze the phospho-aspartyl bond in the phosphorylated response regulator, thus resetting the two-component system^10,11^. Whereas the kinase activity has been extensively studied^12^, the importance of the phosphatase activity has been appreciated more recently^13–15^. In two-component systems, a dynamic balance between kinase and phosphatase activities determines the net output and downstream physiological effects. These different conformational states are necessary for balancing their activities^16,17^.

In contrast to the typical transmembrane HK receptors, light sensitive receptors are frequently soluble. This facilitates their structural and mechanistic analyses^4,18,19^. As a case in point, phytochromes are red/far-red light-sensing photoreceptors that regulate diverse physiological processes in plants, fungi, and bacteria^20^, e.g., chromatic adaptation and phototaxis in prokaryotes^21^. Bacterial phytochromes (BphPs) usually belong to two-component signaling systems, with a cognate response regulator commonly encoded in the same operon^18,22,23^. BphPs comprise an N-terminal photosensory module (PSM) with PAS (Period/ARNT/Single-minded), GAF (cGMP phosphodiesterase/adenylyl cyclase/FhlA) and PHY (Phytochrome-specific) domains^24^. The PSM binds a biliverdin IXα chromophore via a thioether linkage to its conserved cysteine within the PAS domain^25,26^. The PSM is followed by a C-terminal output module, usually a HK domain.

Photoactivation by red and far-red light drives biliverdin *Z/E* isomerization, which underlies the phytochrome switch between its red light-absorbing (Pr) and far-red light-absorbing (Pfr) states^27^. Phytochromes can revert to their dark-adapted resting state thermally in dark. Whereas the Pr state is the dark-adapted and most stable state in canonical phytochromes, it is the Pfr state in bathy phytochromes^28^. As first demonstrated for the model bacteriophytochrome from *Deinococcus radiodurans* (DrBphP), light induces extensive structural changes in the photosensory module that are relayed to the output module^29^.

In terms of enzymatic activity, the dark-adapted Pr state exhibited higher kinase activity than the Pfr state in cyanobacterial phytochrome Cph1,^27,30,31^, similar to other bacteriophytochromes^32^. In particular, the bacteriophytochrome from *Agrobacterium fabrum* (AgP1) displays histidine kinase activity in its resting Pr state^28,33^; in the Pfr state, the autophosphorylation and phosphotransfer reactions are down-regulated by 2-fold and 10-fold, respectively^28^. This kinase activity of the AgP1 has been shown to control bacterial conjugation^34^. By contrast, no kinase activity has been demonstrated for DrBphP, notwithstanding close sequence homology and the elaborate structural changes this receptor undergoes under light^22,29^. Despite the eminent role of DrBphP as a paradigm for photoreception, the enzymatic activity and the physiological role of this model phytochrome have hence remained enigmatic.

Here, we unravel this long-standing puzzle by studying the enzymatic activity and interactions of DrBphP and AgP1, as two canonical bacteriophytochromes with HK effector domains. By pursuing an integrated biochemical and structural strategy, we show that despite close homology, AgP1 acts as a histidine kinase whereas DrBphP functions as a light-activated phosphatase. Our biochemical and structural data pinpoint two key residues proximal to the catalytic histidine that affect the balance between the kinase and phosphatase activities. Together, the two phytochromes provide soluble, light-controllable systems with opposite activities for the study and application of two-component signaling.

## RESULTS

### The spectral properties of DrBphP are affected by DrRR

UV-vis absorption spectroscopy allows to study how the response regulators potentially affect the photoactive states of the bacteriophytochromes. For comparison, we generated a hybrid receptor, denoted as ‘Chimera’, which comprises of the DrBphP PSM and the AgP1 HK domain (Figure 1A). DrBphP, AgP1 and Chimera all showed typical absorption spectra with Soret and Q-band absorption peaks for both states, which were unaffected by the addition of the cognate RR (Figure 1B). The thermal reversion of phytochrome samples after saturating red light (655 nm) exhibited multiple exponential phases in all cases, irrespective the presence of the RR (Supporting Figure S1A). The recovery of AgP1 was faster than that of DrBphP. The thermal reversion of the Chimera was between that of DrBphP and AgP1. Earlier studies indicate that the dark reversion in phytochromes is affected by the dimerization interfaces in both the PSM and HK domain^35^. In line with this notion, the spectral characteristics of the Chimera are governed by both the AgP1 HK and DrBphP PSM.

**FIGURE 1.**
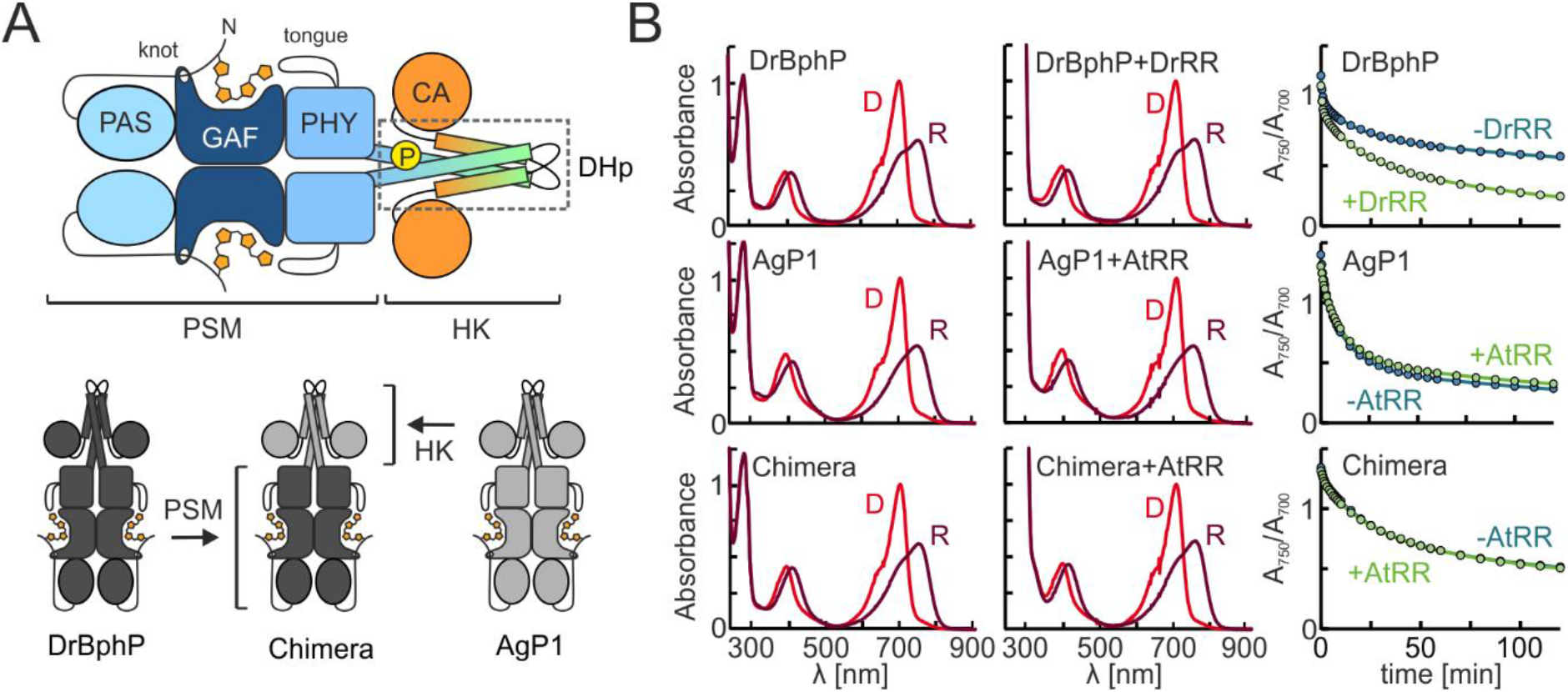
Overall structure and UV-vis spectroscopy of DrBphP, AgP1, and their chimera samples with and without their cognate response regulator. **A**. Schematic representation of a canonical bacteriophytochrome with an effector histidine kinase (HK) domain. The site of phosphorylated histidine is indicated as letter ‘P’. In addition, a schematic presentation of the phytochrome chimera is shown, where the photosensory module (PSM) of DrBphP is combined with the HK domain of AgP1. Domain abbreviations: Period/ARNT/Single-minded (PAS), cGMP phosphodiesterase/adenylyl cyclase/FhlA (GAF), Phytochrome-specific (PHY), Histidine kinase (HK), Dimerization Histidine phosphotransfer domain (DHp), catalytic ATP-binding domain (CA). **B**. The absorption spectra of the BphP HKs with and without RR in dark (D) or under red light (R), and their dark reversion kinetics. The dark reversion is shown as an A_750_/A_700_ ratio over time, where 0 min corresponds to the time the 655 nm illumination ceased. The dark reversion rates of AgP1 and Chimera were unaffected by AtRR, but DrBphP reversion is facilitated by DrRR.

The dark reversion kinetics of AgP1 and Chimera were unaffected by the response regulator from *Agrobacterium fabrum* (AtRR), but that of DrBphP was significantly accelerated by *Deinococcus radiodurans* response regulator (DrRR). This finding indicates that DrRR binds to the DrBphP HK, favoring the Pr state conformation. Interestingly, this contrasts the *Arabidopsis thaliana* phytochrome B, where the binding of the phytochrome-interacting factor (PIF) stabilizes the Pfr state^36^.

### DrBphP interacts with DrRR more strongly than AgP1 with AtRR

To further analyze the interaction between phytochromes and their RRs, we applied size-exclusion chromatography and fluorescently labeled RR proteins (Supporting Figure S1B). Indicative of a binding, the addition of DrBphP caused a shift in the retention of EGFP-labeled DrRR. The interaction was independent on the photoactivation of the protein. By contrast, the EGFP-AtRR retention was not significantly affected by the presence of AgP1, suggesting a weak interaction between AgP1 and AtRR (Supporting Figure S1).

To further investigate the BphP/RR interactions, we resorted to surface plasmon resonance (SPR). The changes in the SPR signal were measured from the response regulator-coupled SPR chips while applying the phytochromes to the sensor surface. The binding of DrRR to DrBphP in the Pr state was evaluated from the steady-state saturation signal (Figure 2A), resulting in a dissociation constant *K*_D_ of (43 ± 8) μM with a 1:1 binding stoichiometry (Figure 2C). This value was verified by Langmuir kinetic analysis which yielded an affinity of comparable strength (*K*_D_ ~ 10 μM). A lower signal amplitude and a *K*_D_ value of (60 ± 7) μM for the DrBphP/DrRR pair was observed upon red-light application. The interaction between AgP1 and AtRR was substantially weaker (*K*_D_ > 200 μM), and no saturation of the binding curve could be reached. The shape of the SPR response graph indicates that the association and dissociation kinetics of the AgP1/AtRR are fast, which precludes the kinetic evaluation. That notwithstanding, the AgP1/AtRR interaction was unaffected by red light (Supporting Figure S2C).

**FIGURE 2.**
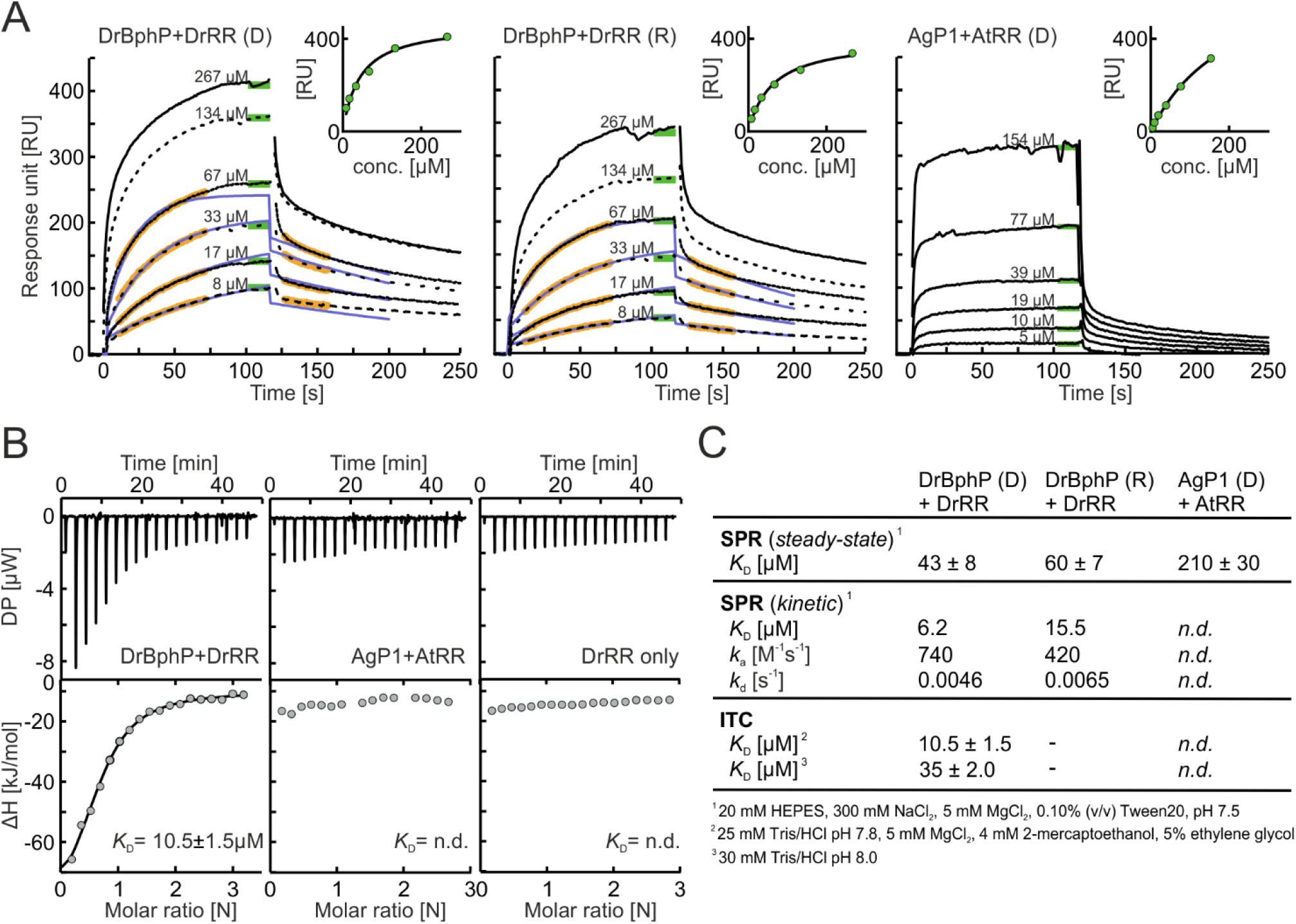
BphP/RR interactions. **A**. Surface plasmon resonance (SPR) measurements. The response regulators were coupled on the sensor surface, and varying concentrations of the corresponding phytochrome were applied on the sensor. The sensorgrams (black lines), the kinetic fits (blue lines) and the parts of the data that were used for kinetic analysis (orange) are indicated. For kinetic analyses, DrBphP concentrations of 8-67 μM were used. Green lines mark the *R_eq_* values that were used for evaluating the steady state affinity data, shown in panel **C**. Inset: Steady-state fit of the concentration series, where *R_eq_* values were used for affinity approximation. DrBphP showed intermediate-affinity binding to DrRR-coupled surface in dark (*K*_D_ = 43 ± 8 μM), which was not notably affected by saturating red light (*K*_D_ = 60 ± 7 μM). Only weak interaction (*K*_D_ > 200 μM) was detected between AgP1 and AtRR. **B**. Isothermal titration calorimetry (ITC) measurements. Differential power (DP) resulting from the injections is plotted against time, and binding enthalpy (ΔH) is plotted against the molar ratio of the proteins. **C**. Table summarizing the results from SPR and ITC analyses. See Supporting Figure S2D for a table with additional SPR fitting values and additional ITC measurements.

As a complementary method, we applied isothermal calorimetry (ITC). The DrBphP/DrRR interaction occurs in 1:1 stoichiometry with a *K*_D_ of (10.5 ± 1.5) μM (Figure 2B), slightly lower than the value attained by SPR (Figure 2C). Unlike in a blue light-regulated HK^37^, this interaction was not affected by the addition of ATP or its analogue AMP-PNP (Supporting Figure S2B). AgP1 binding to AtRR could not be detected by ITC (Figure 2B), consistent with the SEC data and the fleeting binding seen in SPR. The binding parameters were similar in a different buffer condition (Figure 2C, Supporting Figure S2A).

Taken together, the interactions of DrBphP and AgP1 with their cognate response regulators were clearly different. Next we studied whether these differences in the interaction parameters correlate with enzymatic activity, as we speculate that the function of these phytochromes is reflected in their interactions.

### AgP1 functions as a histidine kinase but DrBphP acts as a phosphatase

We characterized the kinase activity of phytochromes by ^32^P-γ-ATP autoradiography (Figure 3A). The autophosphorylation reaction of AgP1 occurred preferably in the Pr state and was reduced under red light illumination, consistent with previous reports^33^. If AtRR was present, it received a phosphate from AgP1 in the phosphotransfer reaction. This reaction occurred preferably in the dark-adapted Pr state but was almost absent under constant red-light illumination (i.e., in Pfr state). This verifies that AgP1 binds to and transfers its phosphate to AtRR in its active Pr state.

**FIGURE 3.**
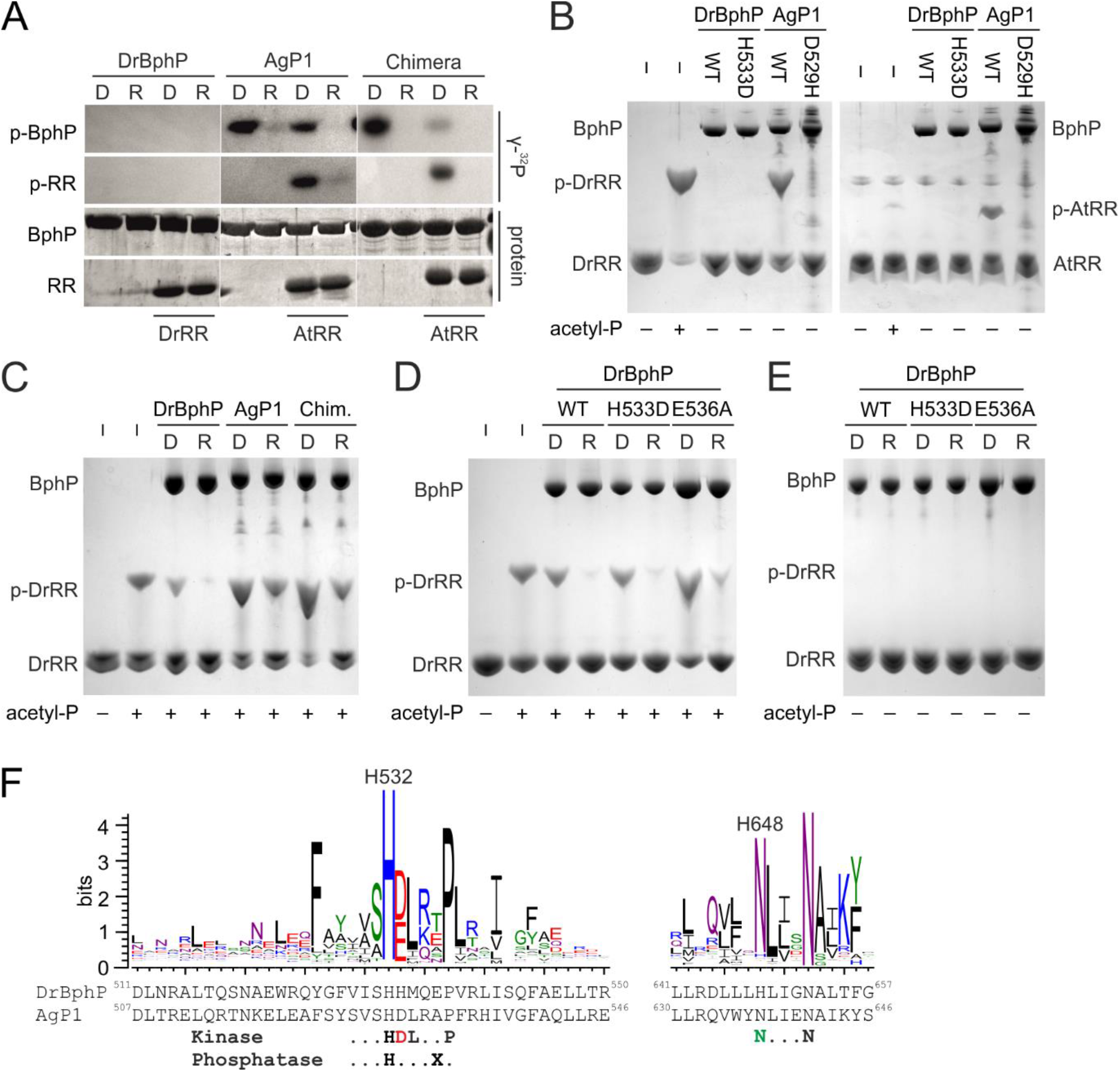
Kinase and phosphatase activity of the phytochromes DrBphP and AgP1. **A**. Kinase and phosphotransfer activity of the phytochromes, detected for radioactive phosphate (γ-^32^P) and total protein. Each phytochrome sample (BphP) was incubated with γ-^32^P-ATP, either with or without the response regulator (RR) protein. DrBphP was incubated with or without DrRR, whereas AgP1 and Chimera were incubated with or without AtRR. Full gel is shown in Supporting Figure S3A. **B**. Kinase activity of DrBphP, AgP1, and their variants with the H+1 residue mutated. Each well was loaded with equal amount of response regulator. The phosphorylated response regulators (denoted p-DrRR and p-AtRR) migrate slower in the gels and are therefore resolved from their unphosphorylated counterparts. **C**. Phos-tag detection of the phosphatase activity of DrBphP, AgP1 and Chimera. Equal amount of phospho-DrRR were applied to each reaction. **D**. Phosphatase activity of DrBphP and its mutants H533D and E536A. Equal amount of phospho-DrRR were applied to each reaction. **E**. All DrBphP variants were inactive kinases. Full gels are shown in Supporting Figure S3. **F**. Sequence logo of 50,000 histidine kinase sequences, shown here for the H box close to the phosphoaccepting histidine (H532 in DrBphP) and the N box in the CA subdomain. The height of each residue name indicates the amount of conservation. The protein sequence of DrBphP and the fingerprint sequence motifs are shown below the graph. In the case of phosphatase activity, the residue at H+4 position varies and is denoted as X.

Intriguingly, DrBphP lacked autokinase or phosphotransfer activities both in the Pr and Pfr state (Figure 3A). The absence of kinase activity is surprising as the DrBphP and its PSM evidently undergo light-induced structural changes that seem to be conserved among other phytochromes^29,38,39^. All homologous bacteriophytochrome HKs studied to date exhibited light-dependent kinase activity. The unusual absence of kinase activity in DrBphP could in principle be due to 1) lack of interaction with the DrRR; 2) inability of its PSM to transduce signals to the HK effector; or 3) inactivity of the DrBphP HK module. Scenario 1 can be ruled out according to the results presented above, which consistently showed interaction between DrBphP and DrRR. To discriminate between the scenarios 2 and 3, the histidine kinase activity of the Chimera was found to function similarly to the wild-type AgP1 with robust autokinase and phosphotransfer activity in the Pr state, but reduced activity in the Pfr state (Figure 3A). This result states that DrBphP undergoes productive structural changes that are conducive to controlling HK activity, thereby ruling out scenario 2.

To address scenario 3, Phos-tag gels were applied where unphosphorylated proteins and their phosphorylated counterparts are resolved based on migration through the gel matrix. In this analysis, unphosphorylated and phosphorylated response regulators were clearly resolved from another (Figure 3B). The assay confirmed that the wild-type AgP1 phosphorylates AtRR preferably in the Pr state and revealed that it cross-phosphorylates DrRR with similar efficiency (Figure 3B). However, like in the radiolabeling assay (Figure 3A), DrBphP lacked kinase activity as it could not produce phosphorylated DrRR (phospho-DrRR).

The residue that immediately follows the catalytic histidine, denoted as H+1, is acidic in the majority of sensor histidine kinases and has been implicated in the autophosphorylation reaction^12^. Whereas AgP1 has Asp529 in this position and thus conforms to the prevalent sequence motif, DrBphP unusually possesses a histidine in the corresponding position 533 (Figure 3F). To test the role of the H+1 position, we generated the AgP1 D529H and DrBphP H533D variants. The D529H mutation rendered AgP1 inactive (Figure 3B), verifying the importance of this acidic residue for the kinase activity. Like the wild-type DrBphP, the H533D variant appeared inactive (Figure 3B and E). Therefore, this single mutation in the H+1 position is not sufficient to rescue the kinase activity of DrBphP.

As sensor histidine kinases may also function as phosphatases^10,40,41^, we tested the DrBphP and AgP1 HKs in that regard. Of particular advantage, the Phos-tag gels allow to assess the dephosphorylation of phospho-RR proteins. To this end, we generated the phosphorylated response regulators chemically by treatment with acetyl phosphate^42^. DrRR was phosphorylated robustly, whereas AtRR responded to the treatment weakly. Phospho-DrRR was then incubated together with ATP and either DrBphP or AgP1. In the reactions, net phosphatase activity would decrease the amount of phospho-DrRR, whereas net kinase activity would increase it. As expected, addition of AgP1 or Chimera led to an increase in phospho-DrRR when incubated in darkness, indicating kinase activity of these proteins (Figure 3C). By contrast, the addition of DrBphP decreased the amount of phospho-DrRR, especially upon red-light exposure. These findings reveal that DrBphP acts as a phosphatase with higher activity in the Pfr state than in the Pr state. The H533D mutation did not unalter the phosphatase activity (Figure 3D).

The residue in the H+4 position is important for phosphatase activity of HisKA family proteins^14^. Indeed, the corresponding residue E356 in DrBphP appeared to have a role in the reaction as its mutation to alanine reduced the phosphatase activity (Figure 3D). Notably, the E356A variant maintained somewhat higher phospho-DrRR amounts not only in the Pfr but also in the Pr state, potentially by shielding the phospho-DrRR from spontaneous hydrolysis during the reaction. Like the H533D and wild-type variants, E536A did not show any histidine kinase activity (Figure 3E). Interestingly, the phosphatase activity in DrBphP depended on the ATP addition, as phospho-DrRR remained unchanged in the absence of ATP (Supporting Figure S3C). This is consistent with the notion that ADP, ATP, or adenosine 5’-[β,γ-imido]triphosphate (AMP-PNP) serves as a cofactor for the phosphatase activity^43^.

### DrRR crystal structure reveals a canonical response regulator dimer

As DrBphP and AgP1 strikingly differ in their enzymatic activity and interactions, we next asked whether these differences can be explained by the structure of the interface between the DHp and RR. To model this interface with confidence, we solved the crystal structure of the response regulator from *D. radiodurans* (DrRR) at 2.1 Å resolution (see Table 1 for crystallographic statistics). The protein crystallized in the tetragonal P41212 space group with four monomers in the asymmetric unit. These monomers form two ‘inverted 4-5-5 dimers’ with a dimerization interface built by a α4–β5–α5 face of each monomer^44^ (Figure 4A), similar to most other phytochrome RR structures with a receiver domain^45–47^. However, the AtRR assumes a ‘hand-in-hand’ dimer^48^, in which the C-terminal extension forms an antiparallel β-strand interface with a sister monomer (Figure 4A). Overall, the structure of the DrRR is highly similar to other reported response regulators. It contains the structural features and the conserved residues critical for its receiver domain function in two-component signaling (Figure 4C). These structural details along with functional results (Figure 3) verify that DrRR can function as a canonical response regulator in a two-component signaling system.

**FIGURE 4.**
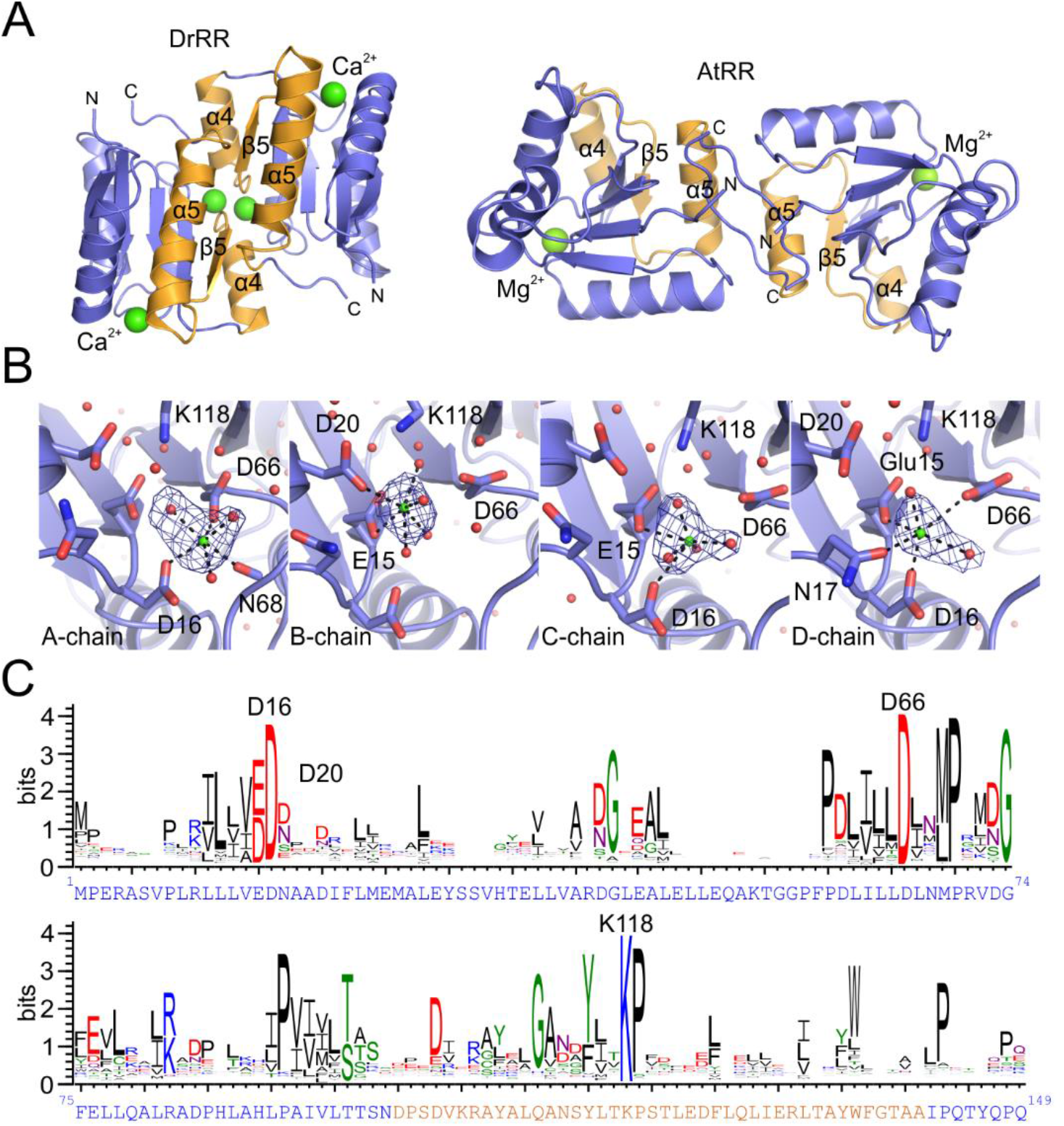
Crystal structure of the *Deinococcus radiodurans* response regulator. **A**. Cartoon representation of the dimeric DrRR and AtRR structures (PDB code 5BRJ for AtRR^48^). The α4–β5–α5 face of each response regulator monomer is in orange, and the rest of the protein is in blue. Ca^2+^ and Mg^2+^ ions at the active sites, N- and C-termini, as well as dimerization helices are marked. In the case of DrRR, a dimer formed by chains A and B is shown. **B**. The active site of DrRR with its Ca^2+^ ions and interacting residues. The localization and its interactions (black dashed lines) of Ca^2+^ ions (green) differ between the chains. The omit difference (Fo–Fc) map of the Ca^2+^ ions are shown as blue mesh at 5.0σ. The omit maps are calculated for each monomer by repeating the final refinement step without the Ca^2+^ ion and the coordinating water molecules. **C**. Sequence logo attained from 50,000 response regulator sequences. The height of each letter indicates the amount of conservation for the corresponding amino acid (one-letter code). The key DrRR residues are shown above the graph, and the full amino acid sequence of DrRR is below the graph. The amino acid sequence is colored as in panel A.

**Table 1.** Crystal data collection and processing statistics.

|  |  |
| --- | --- |
| **Data Collection<sup>a</sup>** |  |
| Space group | P 41 21 2 |
| Cell dimensions |  |
| *a*, *b*, *c* (Å) | 87.65, 87.65, 181.21 |
| $\alpha$, $\beta$, $\gamma$ (°) | 90.00, 90.00, 90.00 |
| Resolution (Å) | 49.74–2.0 (2.15–2.10)<sup>a</sup> |
| *R*<sub>merge</sub> | 0.172 (2.882) |
| CC<sub>1/2</sub> | 0.999 (0.503) |
| *I*/$\sigma$(*I*) | 12.01 (1.09) |
| Completeness (%) | 100.0 (100.0) |
| Redundancy | 4.13 (3.94) |
| Wilson B factor | 49.54 |
| **Refinement** |  |
| Resolution (Å) | 49.74–2.10 (2.15–2.10)<sup>a</sup> |
| No. of reflections | 39,970 (2,891)<sup>a</sup> |
| *R*<sub>work</sub>/*R*<sub>free</sub> | 0.181/0.218<sup>b</sup> (0.330/0.318) |
| Overall B factor | 59.84 |
| No. of atoms |  |
| Protein | 4,389 |
| Heterogen<sup>c</sup> | 9 |
| Water | 240 |
| **Geometry** |  |
| RMSD |  |
| Bond lengths (Å) | 0.013 |
| Bond angles (°) | 1.811 |
| Ramachandran |  |
| Favored (%) | 96 |
| Allowed (%) | 16 |
| Outliers (%) | 5 |
| **PDB Code** | 6XVU |

<sup>a</sup> Outer shell values used in the refinement are in parentheses.
<sup>b</sup> Test set for *R*<sub>free</sub> calculation constitutes 5% of total reflections that were randomly chosen.
<sup>c</sup> This includes nine Ca<sup>2+</sup> atoms.

Notably, the crystal structure of DrRR contained Ca^2+^ instead of Mg^2+^ ions found in other response regulator structures^45–48^. The Ca^2+^ ions played a central role in this crystal form, as their replacement with Mg^2+^ did not allow crystal formation. Ca^2+^ ions occupy the active site of the DrRR in a similar way to Mg^2+^ in the AtRR structure^48^. Although Ca^2+^ is chemically similar to Mg^2+^, its larger size leads to diffuse coordination of the ion in the active sites (Figure 4B) and 45% higher B-factors compared to Mg^2+^ ions modelled at the same sites. Consequently, the Ca^2+^ interactions differ between the four monomers in the asymmetric unit, being most similar to AtRR in monomer A^48^. In each case, the Ca^2+^ ions are hexagonally coordinated to surrounding atoms, which involve water molecules, the side chains of Glu15, Asp16, Asn17, the phospho-accepting Asp66, and the main-chain oxygen of Asn68 (Figure 4B).

### Complex structures show different interactions in DrBphP and AgP1

To analyze how the interplay of the HK and RR proteins impacts on two-component signaling, we prepared models for the DrBphP/DrRR and AgP1/AtRR pairs by utilizing a homologous complex structure^9^ and the crystal structures of the DrRR and AtRR^48^. Support for the physiological relevance of the resultant structural interface also derives from a covariance analysis of cognate HK/RR pairs, as pioneered by Laub and coworkers^49^. In contrast to prior analyses, we focused on a set of hybrid receptors which comprise HK and RR moieties in a single polypeptide chain, thus allowing to assign interacting, cognate HK/RR pairs with high confidence. The multiple alignment of several thousand such receptors revealed strong residue covariation not only within the HK and RR parts but also in between them^50,51^. When mapped on the structural model of the complex, strong pairwise residue covariation localized to the HK/RR interface (Supporting Figure S6), speaking for realistic complex models.

To address the stability of the complex models in solution and to reveal the underlying molecular interactions, we conducted classical MD simulations at 300 K and 1 atm pressure with 0.1 M NaCl concentration using the Gromacs molecular dynamics package^52^. Over a 200 ns trajectory, both the DrBphP/DrRR and AgP1/AtRR complexes were stable. The RMSD equilibration times for protein backbone atoms were around ~60 ns for DrBphP/DrRR complex and ~80 ns for AgP1/AtRR (Supporting Figure S4), suggesting that the interactions are more defined and stronger in the DrBphP/DrRR complex. Starting from the 100 ns time point of the trajectory, we extracted snapshots at 10 ns intervals and performed extended analysis of their interactions. Representative snapshots are shown in Figure 5, all snapshots in (Supporting Figure S4).

**FIGURE 5.**
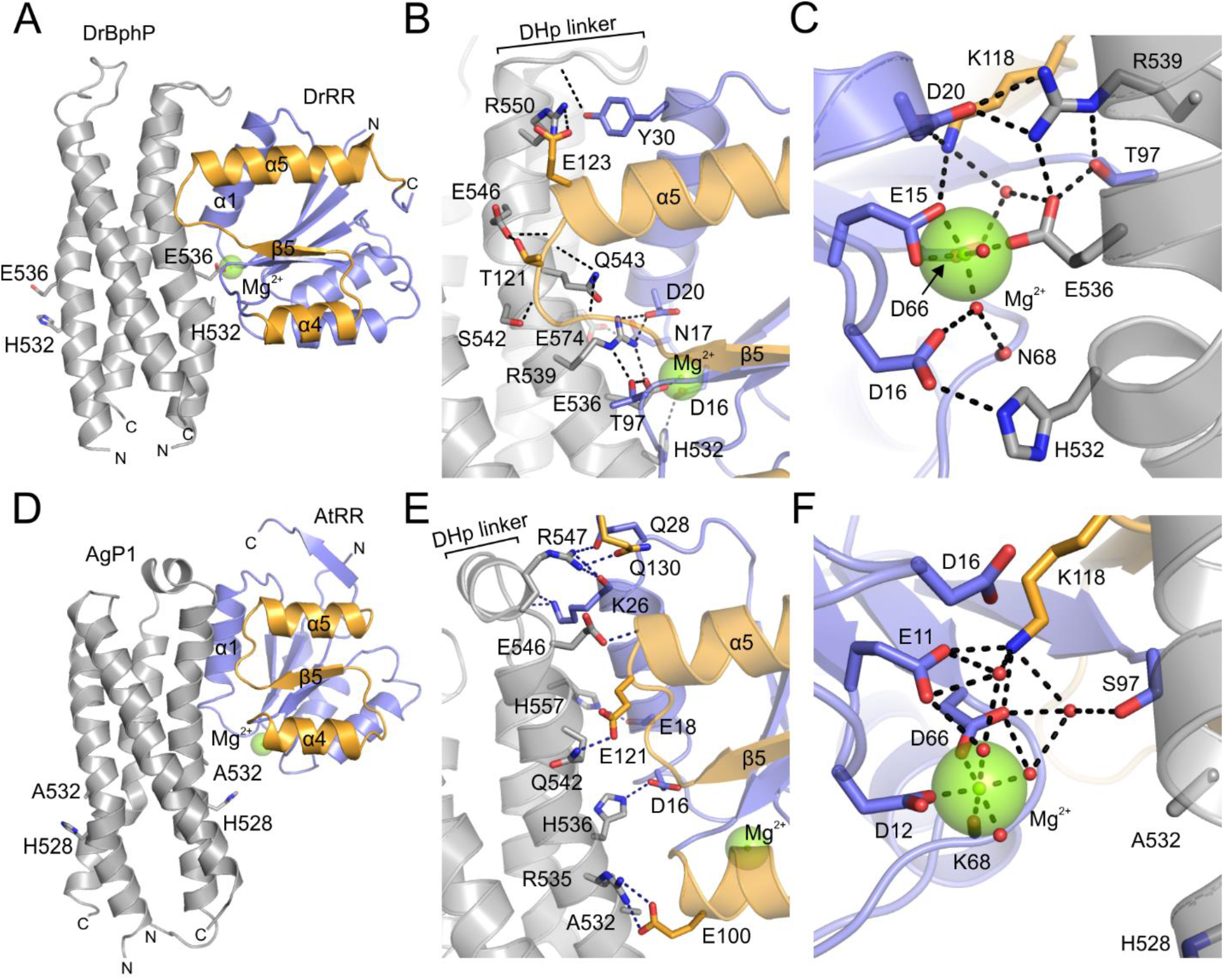
Complex models of response regulators and the interacting DHp domains. **A–C**. Structure and interactions of the DrBphP/DrRR complex, with the overall structure of the complex shown in panel A, detailed interactions in panel B, and the interaction network around the Mg^2+^ in panel C. **D–E**. Structure and interactions of the AgP1/AtRR complex, with the overall structure of the complex shown in panel D, detailed interactions in panel E, and the interaction network around the Mg^2+^ in panel F. Representative structural snapshots from the relaxation simulations are shown. Surrounding water box with Na^+^ and Cl^-^ ions are omitted for clarity reasons, and only one monomer of the response regulator dimer is shown.

Overall, the interactions between AgP1 and AtRR were transient and more variable than the ones in the *D. radiodurans* pair, as gauged by the larger overall RMSD values of the AgP1/AtRR complex and higher mobility of the atoms throughout the MD trajectory (Supporting Figure S4). Analysis of the snapshots in the PISA server^53^ revealed that both complex interfaces have hydrophobic core regions. The average solvation free energy upon formation of the interface indicated that this interface is more extensive in the DrBphP/DrRR complex (−11.3 kcal/M) than in the AgP1/AtRR complex (-5.8 kcal/M).

The RRs interacted mainly through their α1 helix (aa. 18–32 in DrRR) that aligns with the helical bundle of the four DHp helices. In addition to this main interface, DrRR has interactions through a loop region (aa. 119–121) that connect strand β5 and helix α5. Notably, the position of this ‘β5–α5 loop’ and the length of the α5 helix differ between DrRR and AtRR, allowing DrRR more extended interactions with its phytochrome partner. The complexes contain polar interactions and well-defined salt bridges that are more pronounced in the *D. radiodurans* complex (Figure 5). Notably, Mg^2+^ ion coordinates a set of interactions between DrBphP and DrRR (Figure 5C), which are absent in the AgP1/AtRR complex (Figure 5F). We note that in DrBphP/DrRR complex, inter-chain salt bridges are less fluctuating in comparison to the AgP1/AtRR complex (Supporting Figure S5B, D).

In DrBphP, the catalytic residue His532 forms a salt bridge to the DrRR residue Asp16, which orients it close to the DrRR phospho-accepting Asp66. Glu536 coordinates with Mg^2+^ and forms additional interactions with Arg539 and DrRR residue Thr97 (Supporting Figure S5A, B). The corresponding residue in AgP1 is alanine (Ala532), and therefore these interactions are absent in the AgP1/AtRR pair. DrBphP residue Arg539 forms a salt bridge with DrRR residue Asp20 and interacts with Thr97, Arg539 and Glu536. The α5 helix is longer in DrRR than in AtRR, which enables additional contacts between the ‘β5–α5 loop’ region (aa. 119–123) and DrBphP residues Ser542, Asn543, Glu546, and Arg550. This positioning of the ‘β5–α5 loop’ guides the side chain of Arg539 to close proximity to Asp20, thus enabling extensive interactions around these residues (Figure 5C). In the case of AgP1, the corresponding residue Arg535 points away from Asp16 of AtRR (Figure 5E).

Additional interactions that stabilize the DrBphP/DrRR complex are mediated by DrBphP residues Glu574 and Arg550. Tyr30 of DrRR interacts with the main chain of Ala554, thus stabilizing this DHp linker loop between the two DHp helices (Figure 5B). In a similar way, AtRR residue Lys26 interacts with Ala550, Lys554 and Ser555 of AgP1, stabilizing a helical linker between the DHp helices (Figure 5E). The AgP1/AtRR complex is further supported by interactions mediated by the side chains of Gln542, Glu546, Arg547, and H557 (Figure 5E, Supporting Figure S5D).

Taken together, there are three central DrBphP residues that interact with the DrRR active site: His532, Glu536 and Arg539. These residues form a defined interaction network that includes a hexagonally coordinated Mg^2+^ ion (Figure 5C, Supporting Figure S5). The corresponding residues are not interacting with the active site in the AgP1/AtRR complex (Figure 5E), leading to active-site structure that resembles free AtRR^48^.

## DISCUSSION

### The bacteriophytochrome from *D. radiodurans* is a light-activated phosphatase

Bacterial phytochromes commonly act as light-regulated histidine kinases in two-component systems^27^. Here, we introduce novel biochemical and structural insight into the activity of these phytochromes and their interaction with response regulators.

AgP1 acts as a red light-repressed histidine kinase that phosphorylates its cognate response regulator AtRR^33^ and that from *D. radiodurans* (Figure 3A, B). Similar cross-reactivity has been reported for a bacteriophytochrome from *Pseudomonas syringae* (PsBphP) which can phosphorylate DrRR^22^. Therefore, it is possible that AgP1 acts promiscuously and phosphorylates other response regulators in bacteria. By contrast, DrBphP did not show any kinase activity but functions exclusively as a phosphatase (Figure 3). Therefore, DrRR is largely phosphorylated non-enzymatically or by other histidine kinases in bacterial cell under dark conditions. DrBphP can dephosphorylate phospho-DrRR upon red-light exposure and thereby elicit physiological responses. Notably, the precise function of DrBphP has been under debate ever since its role in the control of carotene production was reported^54^. Our results now settle this long-standing debate and show that DrBphP is a biochemically active protein that dephosphorylates the DrRR, rather than phosphorylating it.

The DrBphP PSM fused with the histidine kinase effector of AgP1 (Figure 1A) produced a functional histidine kinase chimera (Figure 3C). This shows that the DrBphP PSM is principally capable of controlling both histidine kinase and phosphatase activities in dependence of light. Consistent with this observation, the well-studied conformational changes of the DrBphP PSM during the Pr-to-Pfr transition^26,29,55^ are generally very similar across various phytochromes^38^, indicating that both histidine kinase and phosphatase modules can be controlled with other phytochrome photosensory modules.

Even prior to the present elucidation of the enzymatic activity of DrBphP, its PSM has provided a versatile building block for novel light-controllable enzymes to be used in optogenetics. Pertinent enzymes have for instance been constructed through fusion of the DrBphP PSM with a cyclic-mononucleotide phosphodiesterase^56,57^, a guanylate/adenylate cyclase^58,59^, and a tyrosine kinase^60^. Our study introduces a red light-regulated HK chimera and phosphatase as a potential addition to the optogenetic toolkit.

### The binding modes of AgP1 and DrBphP support different functionalities

Phytochrome photoactivation entails large-scale structural changes in the photosensory module, which are then relayed to the output module^26,29,55^. Although the molecular details are under debate and may differ between receptors, the conformational changes in the DHp bundle likely include rotation, bending, or changes in register of the constituent helices^18^. These conformational transitions can then change the interactions and/or enzymatic activity of the output HK domain.

The binding of DrBphP and AgP1 to their cognate response regulators differ from each other, which may be integral to their respective activity profile. This difference manifested in dark reversion (Figure 1B), in SEC profiles (Supporting Figure S1), and in SPR and ITC analyses (Figure 2). The binding of AtRR to AgP1 was weak and transient, but the binding of DrBphP to DrRR had moderate affinity (*K*_D_ ~10 μM) and slower association kinetics. Our structural data indicate that the binding interfaces in both complexes are generally similar but differ in their details: The DrBphP/DrRR complex had relatively stable interactions, whereas the interactions appeared transient and less defined in the AgP1/AtRR complex (Figure 5).

We did not detect clear light-induced change in binding in the BphP/RR pairs, which is consistent with structural evidence that the RR binds to the DHp domain in a similar way regardless of whether the receptor resides in the kinase or phosphatase state. Notably, the relevant interaction epitope of the DHp domain experiences only minor structural changes upon HK (in)activation^17,18^. The CA domain on the other hand binds to a DHp region that varies upon light activation. This may lead to a change in kinase activity, either through a difference in CA binding, through varying the accessibility of the catalytic histidine^17^, or both. The binding sites of the CA domain and RR partially overlap, as also visible in the covariance analyses of the interfaces (Supporting Figure S8), which should create competition between the two binding schemes.

The structural changes in the DHp domain may facilitate the switch between the CA binding during the autokinase reaction and the RR binding during the phosphotransfer or phosphatase reaction. As the RR competes with the CA domain for binding to the DHp domain, transient interactions between the molecules would be favored in the kinase reactions. Structural asymmetry, observed for several HKs in their kinase-active state, may also facilitate the alternating binding of CA and RR^4,18,61^. Although the phosphatase reaction is greatly facilitated by a CA domain^43^ and ATP (Supporting Figure S3C), the CA binding during the phosphatase reaction likely differs from that of kinase reaction. This difference may allow relatively slow binding kinetics between the DHp bundle and the phosphate-presenting response regulator.

### Two residues in the DHp helix govern the HK activity

Two residues within the DHp domain, at positions +1 and +4 relative to the active-site histidine, have been implicated as particularly important for the enzymatic activity in the HisKA family. First, the autophosphorylation reaction involves a nucleophilic attack by the histidine to the γ-phosphate of ATP. This is facilitated by an acidic residue (aspartate or glutamate) in the H+1 position acting as a general base^12^. Second, a threonine or asparagine residue in the H+4 position governs phosphatase activity^14^. There is no crosstalk between the residues, as the H+1 position does not contribute to the phosphatase activity and H+4 position is not required for the kinase activity^14^.

A large-scale sequence analysis of histidine kinases shows that the acidic residue in the H+1 position is strictly conserved among the HisKA family (Figure 3F), underlining its importance. If this residue is mutated, the kinase activity is impaired, as indicated by our results on the D529H mutant of AgP1 and wild-type DrBphP (Figure 3B). A histidine in the H+1 position is very rare among the HK sequences (Supporting Figure S7). Although important, the activity of DrBphP could not be rescued only by introducing an aspartate to this H+1 position (Figure 3B, E). In HisKA proteins, the acidic residue in H+1 is accompanied by an asparagine in the N-box of the CA domain, which stabilizes the active HK conformation and participates in phosphoryl transfer from ATP to the catalytic histidine^40^. Indeed, this asparagine shows a high level of conservation within the HisKA family (Figure 3F). Our model of AgP1 HK supports this interaction between H+1 aspartate (Asp529) and the N-box asparagine (Asn637) (Figure 6A). By contrast, this interaction is likely absent in DrBphP, as both these residues are histidines (His533 and His648) (Figure 6A). Consistent with this observation, the sequence analysis indicates that when the H+1 position is a histidine, the conservation of the N-box asparagine is lost (Supporting Figure S7A).

**FIGURE 6.**
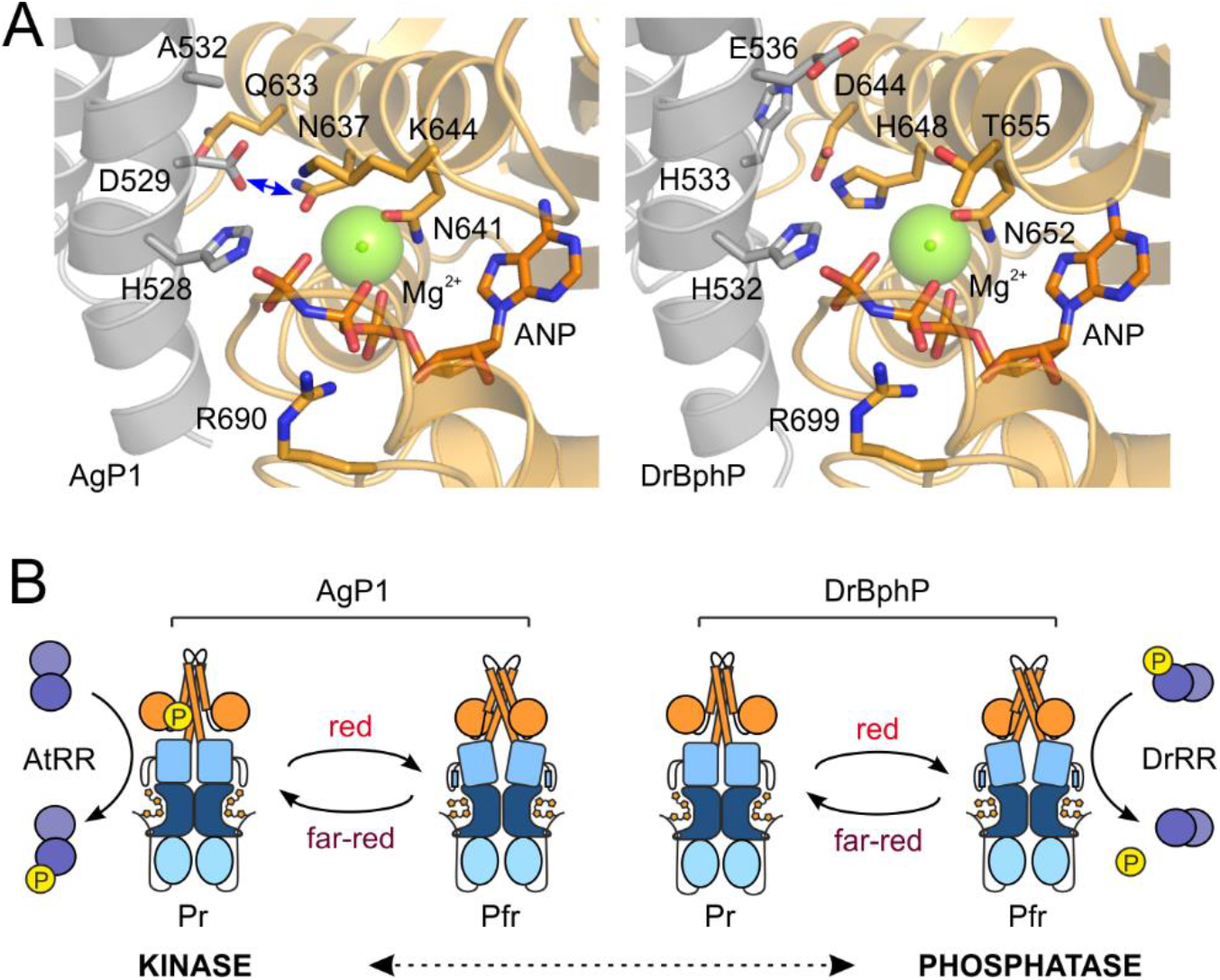
Structural comparison of residues important for the phosphotransfer reaction. **A.** Homology models of AgP1 and DrBphP that are based on the histidine kinase domain structure of HK853/EnvZ chimera in its phosphotransfer state (PDB 4KP4)^12^. The central residues are denoted as sticks, dimerization and phosphohistidine (DHp) domain is shown in grey, and the catalytic ATP-binding (CA) domain is shown in orange. In AgP1, the potential interaction between His528 and Asn637 is show as blue double-headed arrow. **B.** Proposed kinase and phosphatase activities of AgP1 and DrBphP. Many histidine kinases can function as both kinases and phosphatases, which may be governed by their DHp orientations^17^. This transition between the two states is illustrated here as vertical line. In the case of the two phytochromes studied here, red light illumination causes conformation that favors phosphatase activity. In resting (Pr) state, the HK conformation favors kinase activity.

DrBphP has Glu536 in the H+4 position, indicating that this residue is central for the phosphatase activity. As previous studies on the phosphatase activity in HisKA proteins have mainly concentrated on threonine and asparagine residues, not much is known about the role of glutamate at this position^14,62–65^. That notwithstanding, glutamate is as conserved at this site as threonine or asparagine (Figure 3F), which implies that all these residues play equally important roles, potentially fine-tuning the extent of the phosphatase reaction. Changing this residue to alanine diminishes or removes the phosphatase activity^14,62,64,65^, like demonstrated presently in the E536A variant of DrBphP and the wild-type AgP1 (Figure 3D).

In our DrBphP/DrRR complex model, Glu536 forms a distinctive interaction with Mg^2+^ in the active site (Figure 5C). As this site would be occupied by the phosphate moiety of the phospho-DrRR, it is likely that Glu536 facilitates the dephosphorylation reaction. Accordingly, the side-chain hydroxyl group of the threonine or asparagine at the H+4 residue may retain similar interactions as the carboxyl group of the glutamate. In addition to Glu536, Arg539 in the H+7 position participates in the interactions at the DrRR active site (Figure 5C), suggesting a subsidiary role in the phosphatase reaction. This is supported by similar conservation patterns of the H+7 arginine and the H+4 glutamate in HisKA proteins (Supporting Figure S7A).

### Model for bacteriophytochrome photoactivation

To conclude, Asp529 in the H+1 position enables AgP1 to function as a histidine kinase in the Pr state. In the case of DrBphP, His533 at H+1 (along with other residues) renders it inactive as a Pr-state kinase, whereas Glu536 at the H+4 position makes it an effective phosphatase in the Pfr state (Figure 3). Thus, the Pr-like conformation of the bacteriophytochromes in general supplies a structural framework for kinase activity, whereas Pfr-like conformation shifts the output activity to a phosphatase (Figure 6B). Indeed, many HisKA family proteins function as both kinases and phosphatases, and these modes of action can be switched by a change of the relative orientation of DHp helices^17^. As demonstrated in the related histidine kinase YF1, blue light prompts quaternary transitions that channel into a register shift and supercoiling of the DHp helices^66,67^. Both kinase and phosphatase activities are therefore supported by changes within the same structural framework. The bidirectional activity would also require both sets of activity-determining residues in the HK domain, which seems to be the case in many phytochromes (Supporting Figure S7B).

Phytochrome function includes several structural tiers that range from the chromophore surroundings to large-scale structural changes in the entire protein. These tiers are in dynamic equilibrium, which can be shifted by the other tiers and by external factors^68^. The level of phytochrome output activity can be considered to be in an equilibrium between histidine kinase and phosphatase activities^18,61^ (Figure 6B). This equilibrium can be shifted to one direction by the light-induced changes in the photosensory module, and tuned by the sequence variation in the HK domain. In this study, we have shown how small differences in sequence dictate opposing enzymatic activities in two canonical phytochromes. In both cases, light controls their enzymatic activity.

## METHODS

### Cloning and DNA material

The phytochrome from *Deinococcus radiodurans* strain R1 (DrBphP, gene DR_A0050) in pET21b(+) plasmid (Novagen) was kind gift from Prof. Richard Vierstra^24,54^, and phytochrome 1 from *Agrobacterium fabrum* strain C58 (AgP1, gene Atu1990) in pQE12 (Qiagen) was a kind gift from Prof. Tilman Lamparter^33^. AgP1 has a spontaneous R603C mutation, which resides on the surface of the CA domain. The mutations to DrBphP (H533D, E536A) and for AgP1 (D529H) were introduced with QuikChange Lightning Multi Site-Directed Mutagenesis Kit (Agilent Technologies). For cloning the chimera construct, DrBphP residues 513–755 were replaced with Agp1 residues 509–745. First, an XhoI restriction site was introduced after DrBphP residue 513 with QuikChange Lightning Multi Site-Directed Mutagenesis Kit (Agilent Technologies). Then, the C-terminal Agp1 fragment (aa 511–755) was ligated between the new XhoI site and an XhoI site right before the C-terminal His6-tag. After introduction of the Agp1 fragment, the new XhoI site was changed to Agp1 residues 509–510 by site-directed mutagenesis. The response regulators from *Deinococcus radiodurans* strain R1 (DrRR, gene DR_A0049) and *Agrobacterium fabrum* strain C58 (AtRR, gene Atu1989)^22^ were produced as a service (Invitrogen). The response regulator constructs were cloned into pET21b(+) vectors (Novagen) by using restriction sites BamHI and XhoI. The EGFP-RR constructs were prepared with Gibson assembly cloning, in which N-terminal T7 tag was replaced with an EGFP-C1 sequence^69^. In addition, a linker of 10 residues (DSAGSAGSAG) was introduced with primers between the RR and EGFP sequences.

### Sample expression and purification

All DrBphP variants and the response regulators were expressed in *Escherichia coli* strain BL21 (DE3) overnight at 20–24 °C, and purified with HisTrap affinity purification followed by size-exclusion chromatography like previously described^70^. The His6-tagged phytochrome samples were first purified with NiNTA affinity purification using HisTrap™ columns (GE Healthcare), followed by size-exclusion chromatography (HiLoad™ 26/600 Superdex™ 200 pg, GE Healthcare) in buffer (30 mM Tris, pH 8.0). No external biliverdin was added to the cell lysate in response regulator purifications. AgP1 and its D529H mutant were expressed in NEB Express^®^ I^q^ *E. coli* strain (New England Biolabs). The purification protocol was identical to other samples with a couple of exceptions: Protease inhibitor mix (ROCHE) and 0.5 mM TCEP were included in the sample before lysis, and affinity purification was conducted in (30 mM Tris/HCl, 150 mM NaCl, 1 mM TCEP) and varying imidazole concentration (5–500 mM). All purified protein samples were concentrated to 25–30 mg/ml in (30 mM Tris/HCl, pH 8.0) and flash-frozen.

### Absorption spectroscopy

The dark reversion of the phytochromes was measured by the absorption spectroscopy using Agilent Cary 8454 UV-Visible spectrophotometer (Agilent). Absorption spectra in the wavelength range of 690–850 nm were recorded from the mixture of response regulator and Pfr-populated BphP sample. The BphP samples were first diluted to 1.0 μM in (25 mM Tris/HCl, pH 7.8, 5 mM MgCl_2_, 4 mM 2-mercaptoethanol, 5% ethylene glycol) to obtain an approximate A_700_ value of 0.1 cm^-1^. Ten times concentration (100 μM) of cognate response regulator was added into the BphP sample. Then the phytochromes were driven to a maximum population of the Pfr state by saturating illumination with 665 nm LED for 3 min, followed by immediate data acquisition in dark. Dark reversion data were recorded at 1 min intervals for first 10 min, which was followed by intervals of 5 min up to 1 hour and finally 10 min intervals until 2 hours. All measurements were performed in dark at ambient conditions (room temperature). The steady-state spectra of Pr- and Pfr-state samples, in presence or in absence of cognate response regulator, were measured in the same buffer as for dark reversion. Pr state spectra were measured from the dark-adapted samples while Pfr spectra were measured after 3 min illumination with 665 nm LED.

The exponential fits from dark reversion data were calculated with Matlab (MathWorks inc.) using Equation 1. In the case of DrBphP samples, three components were used for fitting, whereas two components were adequate for the rest of the samples.

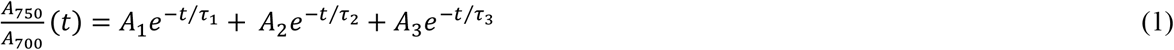

Where *t* is time, *A_700_* and *A_750_* are absorption values at specified wavelength, *A_n_* is the decay amplitude of the absorbance-ratio, and *τ_n_* the time constant of the decay component.

### Size-exclusion chromatography (SEC) with multiwavelength detection

The size-exclusion measurements (Supporting Figure S1) were conducted as previously described^71^. For sample separation, a Nanofilm SEC-250 (300 mm × 4.5 mm) column (Sepax Technologies, Delaware, US) was used with (25 mM Tris/HCl pH 7.8, 5 mM MgCl_2_, 4 mM 2-mercaptoethanol, 5% ethylene glycol) as a mobile phase. Experiments were executed with 350 μL/min flow-rate using the Shimadzu HPLC VP10 pumping system (Shimadzu Corporation, Kyoto, Japan). The eluant was detected at 1.5625 Hz with a diode array UV-Vis detector (SPD-M10A, Shimadzu) and at 2.00 Hz with a fluorescence detector (RF-10A_XL_, Shimadzu). For each run, 20 μL of sample mixture was injected briefly after pre-illumination with 785 nm or 655 nm light. The protein concentrations used were 6 μM (EGFP-DrRR), 30–50 μM (DrBphP), AgP1 60 μM (AgP1), and 1 μM (EGFP-AtRR). To exclude spectral overlap between the samples, 700 nm absorption was used for BphP detection and 488/509 nm excitation/emission was used for EGFP-RR detection. The Gel Filtration standard (Bio-Rad) was used according to the manufacturer’s instructions. The molecular weight estimates were determined by calculating a standard curve of marker proteins Vitamin B12 (1.35 kDa) myoglobin (17 kDa), ovalbumin (44 kDa), γ-globulin (158 kDa), and thyroglobulin (670 kDa).

### Surface plasmon resonance (SPR)

For surface plasmon resonance measurements, phytochrome samples were dialyzed overnight to (20 mM HEPES, 300 mM NaCl_2_, 5 mM MgCl_2_, 0.10% (v/v) Tween20, pH 7.5) with a Spectra/Por^®^ Micro Float-A-Lyzer Dialysis Device (Spectrum Laboratories, USA). The measurements performed using Biacore X instrument (GE Healthcare). Response regulators were coupled onto carboxymethyldextran hydrogel-coated SPR Sensorchip (XanTec bioanalytics GmbH) according to manufacturer instructions. Each response regulator was coupled onto chip surface as 3 mg/mL (150 μM) in an acetate buffer (20 mM sodium acetate, pH 4.2) using EDC/NHS coupling protocol. The remaining activated groups on the sensor chip were then quenched with (1 M ethanol-amine-HCl, pH 8.5). The measurements were conducted by injecting 40 μL phytochrome sample at 20 μL/min, followed by wash step with (20 mM HEPES, 300 mM NaCl_2_, 5 mM MgCl_2_, 0.10% (v/v) Tween20, pH 7.5). Samples were either pre-illuminated with far-red (785 nm) or red (655 nm) LED light before injection, and all measurements were done under dim light at room temperature.

The sensorgrams were analyzed using the BIAevaluation-software version 4.1 (Biacore Life Sciences). The sharp peaks corresponding to the injection start (0 s) and stop (120 s) in each sensorgram were excluded from the analysis. For kinetic fit, a simple 1:1 interaction model between analyte and immobilized ligand was applied, followed by simultaneous fit of k_a_/k_d_ kinetics. The model is equivalent to the Langmuir isotherm for absorption to a surface. Steady state binding levels (*R*_eq_) were obtained by fitting a horizontal straight line to a chosen section of the sensorgrams (blue lines in Figure 5) and determining the average response. *R*_eq_ values (y) and concentrations (x) were plotted in Origin 2018b and a nonlinear simple fit was obtained using the following Equation 2 where A stands for concentration at *R*_qeq_.

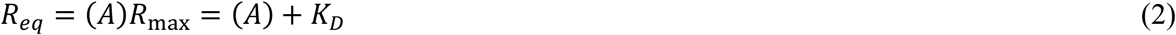

### Isothermal calorimetry (ITC)

Isothermal calorimetry was conducted with MicroCal PEAQ-ITC (Mavern Pananalytical, United Kindom). For the measurements, the purified protein (in 30 mM Tris/HCl pH 8.0) were diluted 1:1 with 2x (50 mM Tris/HCl pH 7.8, 10 mM MgCl_2_, 8 mM 2-mercaptoethanol, 10% ethylene glycol). BphP (30–50 μM, 300 μL) was loaded in the sample cell and RR (750–800 μM, 75 μL) was loaded in the injection syringe. To verify the Pr state of the BphP samples, they were briefly illuminated with 785 nm LED light just before sample application to the cell. The system was equilibrated to 25 °C with a stirring speed of 750 rpm in dark. Injection scheme started with a 0.4 μL response regulator injection, followed by 2 μL injections every 150 s. The ITC measurement in (30mM Tris/HCl, pH 8.0) were made using a Micro-200 ITC (MicroCal, Malvern). The concentrations used were 170–250 μM (BphP) and 750–800 μM (RR). BphP sample (206 μL) was loaded into the sample cell and RR (70 μL) was loaded into the injection syringe. The system was equilibrated to 25 °C with a stirring speed of 750 rpm. The injection scheme started with a 0.2 μL injection followed by 2 μL injections every 180 s. In both measurements, background signal was estimated by injection of response regulator into buffer and buffer into phytochrome with the same parameters. All data from triplicate experiments were analyzed using ORIGIN 7-based MicroCal PEAQ-ITC Analysis Software (Malvern Panalytical). The curves were fitted into a single-site binding isotherm with the first injection excluded. The *K*_D_ value was reported as ± SD from three repeats.

### Radiolabeled kinase assay

The radiolabeled kinase assay was done in a similar way to Lamparter et al.^33,72^. Purified BphPs and RRs were diluted to approximate concentrations of 3.5 μM (0.3 mg/ml) and 9 μM (1.7 mg/ml), respectively, in (25 mM Tris/HCl pH 7.8, 5 mM MgCl_2_, 4 mM 2-mercaptoethanol, 5% ethylene glycol), and pre-illuminated briefly with saturating 785 nm LED light. Reaction was started by adding 3.7 kBq of [γ-^32^P]ATP (PerkinElmer) in a total reaction volume of 10 μL. The samples were then incubated at 25 °C either in dark or under constant 655 nm LED illumination (5 mW/cm^2^) for 20 min. The reaction was stopped by adding SDS sample buffer. The samples were then separated on 12% SDS-PAGE, and the gels were stained with Serva Blue, followed by drying in vacuum drier. The dry gels were then photographed and their radioactivity was monitored with an X-ray film. The experiment was repeated three times.

### Protein phosphorylation by Acetyl phosphate and Phos-Tag detection

In order to create phosphorylated response regulator we adapted the method described by McCleary and Stock^42^. There, response regulators (2–3 μg) were incubated with 50 mM acetyl phosphate for 30 min. The reactions were conducted at 37 °C in (25 mM Tris/HCl pH 7.8, 5 mM MgCl_2_, 4 mM 2-mercaptoethanol, 5% ethylene glycol), followed by buffer exchange to (30 mM Tris/HCl, pH 8.0) with Vivaspin centrifugal concentrator (Sartorius, Germany). The final phosphoprotein concentrations were adjusted to 1.5 mg/ml (80 μM). Both kinase and phosphatase reactions were conducted in (25 mM Tris/HCl pH 7.8, 5 mM MgCl_2_, 4 mM 2-mercaptoethanol, 5% ethylene glycol), where all the desired proteins (2–4 μg each) were incubated in 10 μl total volume at 25 °C, with or without 1mM ATP. The reactions were started by adding ATP to the mixture and incubated either in dark or under saturating 657 nm red light. After 20–30 min, the reactions were stopped by adding 5x SDS loading buffer. For the mobility shift detection of phosphorylated proteins, we applied Zn^2+^-Phos-tag^®^ SDS-PAGE assay (Wako Chemicals). The 9% SDS-PAGE gels containing 25-μM Phostag acrylamide were prepared, and 10 μl of each reaction were run at 40 mA/gel at room temperature according to manufacturer instructions.

### Crystallography

DrRR was crystallized with hanging drops vapor diffusion method. The protein of 10 mg/ml concentration was mixed in a 1:1 ratio with reservoir (0.1 M HEPES pH 7.5, 0.3 M CaCl_2_, 25% PEG400). Crystals formed in few days and were flash-frozen in the reservoir solution containing 15% glycerol. The diffraction data were collected with 0.873 nm wavelength in beamline ID23-2 at the European Synchrotron Radiation Facility (ESRF). The data were processed with the XDS program package version January 26, 2018^73^. The crystals belonged to space group P41212 with two dimers in an asymmetric unit. The initial phases were solved by molecular replacement with Phaser^74^ As for a search model, a DrRR homology model was produced on-line with SWISS-MODEL workspace^23,75^ and a crystal structure of a cyanobacterial response regulator RcpA (PDB code 1K68) as a template^47^. The structure was further refined with REFMAC version 5.8.0135^76^ with automatic weighting and automatically generated local NCS restraints. The model building was done with Coot 0.8.2.^77^. For the final refinement cycles, six TLS regions for each protein chain were implemented from the TLS Motion Determination (version 1.4.0) web server^78^. The final structure had R_work_/R_free_ of 0.181/0.218. Statistics from data collection and refinement can be found in Table 1. Figures from crystal structures and complex models were created with the PyMOL Molecular Graphics System version 2.3.3 (Schrödinger, LLC).

### Computational modelling

For computational simulations, DrBphP/DrRR and AgP1/AtRR complexes were constructed based on a crystal structure containing a sensor histidine kinase and its response regulator from *Thermotoga maritima* (PDB: 3DGE)^9^. Homology models consisting the dimeric DHp bundle of DrBphP (aa 520–592) and AgP1 (aa 513–584) were created on-line with SWISS-MODEL workspace^23,75^ by using the corresponding DHp part of the *T. maritima* histidine kinase (aa 248–316) as a template structure^9^. As for the response regulators, the crystal structures AtRR response regulator (PDB: 5BRJ)^48^ and DrRR (this paper) were applied as dimers. Waters that clashed with the interface and the phosphates at the active sites were not included in the models, whereas the Ca^2+^ and Mg^2+^ ions from the response regulator structures were left intact.

Gromacs 2018.8^52^ classical molecular dynamics package has been used to perform further modelling and simulations. Both AgP1/AtRR and DrBphP/DrRR complexes has been converted into the Gromacs topology, solvated within 15×15×15 nm periodic cubic box of water and neutralized with counterions. In case of DrBphP/DrRR complex we have replaced Ca^2+^ ions which resides in DrRR crystal structure with Mg^2+^ to be consistent with kinetic studies in solution. Additional Na^+^ Cl^-^ ions has been added to the neutralized cell in order to achieve 0.1M total concentration of salt. Amber03^79^ forcefield has been used for the proteins while water has been modelled with TIP3P^80^ parameters.

Classical molecular dynamics simulations have been performed using the following protocol: at first we have minimized our systems for 10000 steps with steepest descent method. Then 200 ns of Classical MD simulation has been performed within NVP ensemble at 300K temperature using a V-rescale thermostat with 0.5 ps time constant^81^ and at 1 atm pressure using Parrinello-Rahman barostat with 1 ps time constant^82^. All bond lengths have been constrained to their equilibrium values, taken from the force field parameters with LINCS method^83^, which allowed us to use 0.2fs time-step for the trajectory integration. A particle mesh Ewald (PME) method^84^ has been used to account for periodic electrostatic interactions with real-space cutoff of 1.5 nm, while Lennard-Jones non-bonded interaction has been treated with a cut-off scheme using a range of 1.5 nm. RMSD of the backbone and RMSF of all the atoms of the proteins with respect to the initial configuration have been extracted from the trajectory.

Starting at 100 ns we have extracted snapshots each 10 ns and performed an extended analysis of interaction between kinases and response regulators. The interactions within the complex structures were analyzed with ‘Protein interfaces, surfaces and assemblies’ (PISA) service at the European Bioinformatics Institute (http://www.ebi.ac.uk/pdbe/prot_int/pistart.html)^53^. In addition, most prominent contacts have been analyzed by plotting contact distances throughout the MD simulation using the standard Gromacs trajectory tools.

### Sequence analysis

To analyze sequence conservation and covariance in sensor histidine kinases, we conducted a BLAST search for the DHp and CA domains (residues 511–755) of DrBphP (Uniprot id BPHY_DEIRA, WP_010889310.1) against the non-redundant (nr) protein sequence database. Using the Biopython interface^85^ and custom Python scripts^86^, we retrieved the top 50,000 sequence hits, corresponding to an *E*-value cutoff of 8.1 × 10^−15^. The sequences were clustered at a 50% identity level with UCLUST^87^, and the 5,404 cluster centroid sequences were determined. The original search sequence (WP_010889310.1) was added, and the sequences were aligned using MUSCLE^88^. The consensus sequence of the alignment, mapped onto the search sequence, was plotted with WebLogo v3.6.0^89^. Based on the alignment, covariance analysis was conducted with PSICOV^51^ as described before^50^. Using custom Python scripts, the score matrix was plotted, and pairwise scores above a cutoff of 0.6 were plotted onto a homology model of the DHp and CA domains of DrBphP (Figure 6A, Supporting Figure S6D). Homology models of DrBphP and AgP1 HK were calculated using SWISS-MODEL^90^ based on the HK853/EnvZ chimera in its phosphotransfer state (PDB 4KP4)^12^.

The sequence analysis of the response regulator proteins was carried out similarly. A BLAST search for the sequence of *D. radiodurans* RR (Q9RZA5_DEIRA, WP_010889309.1) provided 50,000 sequences with an *E*-value cutoff of 2.5 × 10^-10^. Clustering at 50% identity yielded 4,338 sequences, to which were added those of the DrRR (WP_010889309.1) and AtRR proteins (Q7CY46_AGRFC, WP_121650967.1). Sequence alignment and logo representation were done as for the histidine kinase data.

For the analysis of covariance between the histidine kinase and the RR (Supporting Figure S6), the above BLAST hits were scanned for proteins containing consecutive DHp, CA and RR domains in a single polypetide chain. To be included in the subsequent analysis, entries were considered if they contained the Pfam HISKA, HISKA_2 or HISKA_3 domains^91^, immediately followed by HATPase_c and Response_reg domains, with each domain not separated by more than 50 residues at maximum. The resultant 6,805 sequences were clustered at 50% identity, which left 5,386 centroid sequences. The amino acid sequences of the DrBphP (residues 511–755) and DrRR were concatenated and added. All sequences were then aligned as above and analyzed by PSICOV^51^. Pairwise scores above a cutoff of 0.6 were plotted onto a structural model of the DrBphP/DrRR complex (Supporting Figure S6C). In a control run, the aligned sequences were split into their DHp/CA and RR parts and randomly recombined before the analysis by PSICOV. The scrambling of the alignment abolished covariation between the DHp/CA and RR parts (Supporting Figure S6A–B).

## ACCESSION NUMBERS

The Protein Data Bank accession number for the structure reported here is 6XVU.

## ACKNOWLEDGEMENTS

This work was supported by Academy of Finland grants 285461 and 296135 (H. T. and J.A.I., respectively), Jane and Aatos Erkko foundation (J.A.I.), Three-year grant 2018–2020 from the University of Helsinki (E.M and H.T.). S.W. and W.W. acknowledge the European Research Council for support (grant number: 279944), and B.S-B. acknowledges Swiss National Science Foundation (P2ZHP2_164991). D.M. acknowledge the BioExcel CoE (www.bioexcel.eu), funded by the European Union contracts H2020-INFRAEDI-02-2018-823830, H2020-EINFRA-2015-1-675728. We acknowledge the European Synchrotron Radiation Facility (ESRF) for providing synchrotron access for crystal data collection and infrastructure support from the Biocenter Finland and the CSC-IT Finnish center for scientific computing for providing computational resources.

We thank Dr. Harald Janovjak (Monash University) for advice on sequence analysis. We also thank M.Sc. Alli Liukkonen (University of Jyväskylä) for the assistance in laboratory, M.Sc. Moona Kurttila and Dr. Jessica Rumfeldt (University of Jyväskylä) for the help in the UV-vis data analysis, and Prof. Jari Ylänne (University of Jyväskylä) for the help with crystallography data and with radiolabeling assay and Prof. Gerrit Groenhof (University of Jyväskylä) for the help with the MD simulations.

## SUPPLEMENTAL DATA

Supplemental Data include seven Supporting Figures S1–7.

**Supporting Figure S1.**
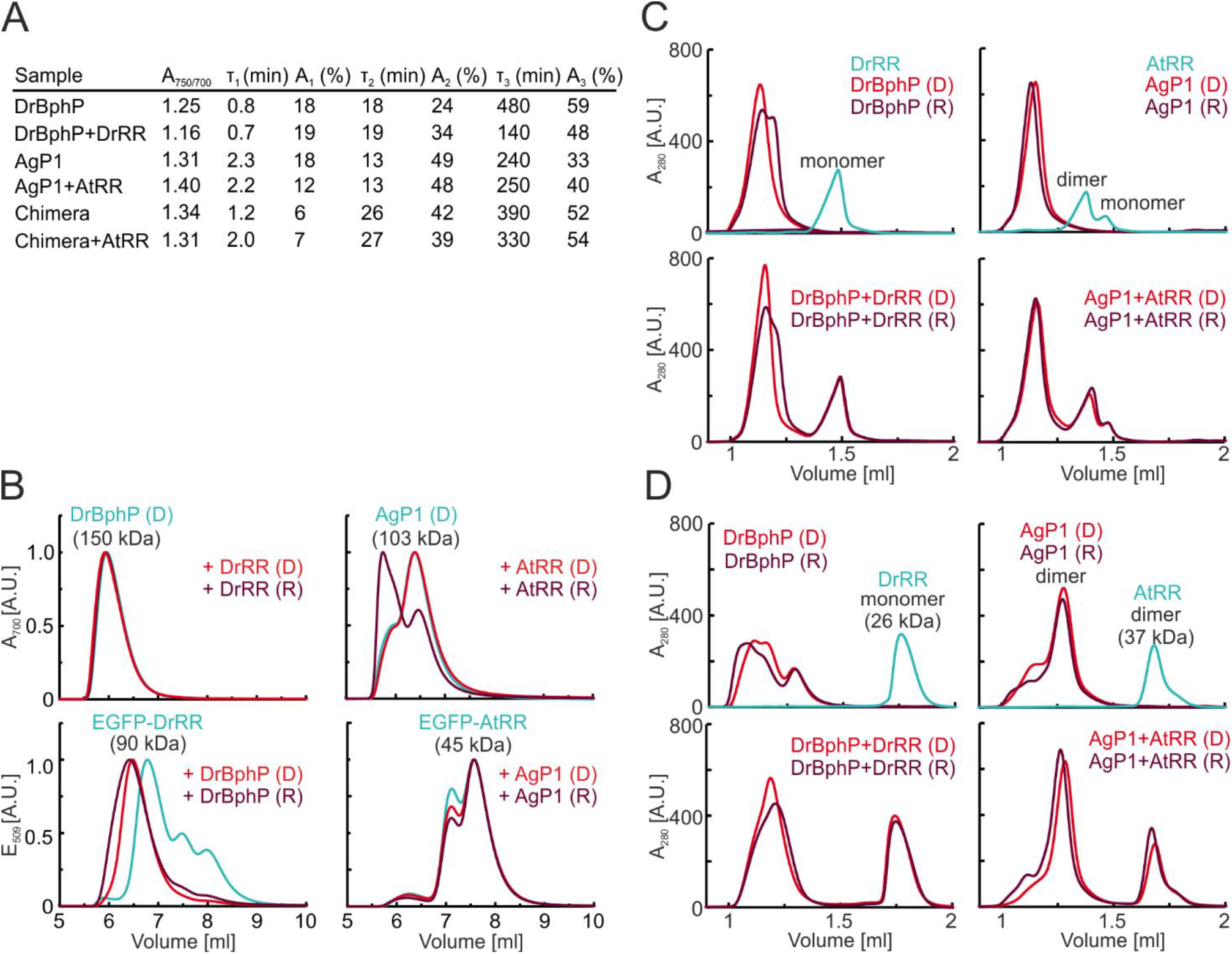
Evaluation of dark reversion data and full gels for radiolabeled kinase assay and size-exclusion chromatography of BphP/RR pairs. **A**. The dark reversion data (Figure 1B) were approximated with an exponential fits. Table summarizes the time constants (τ_n_) and decay amplitudes (A_n_) of the reversion with and without response regulator. Satisfactory fits were obtained by using a sum of three decay components, resulting root mean square error (RMSE) values of 0.003–0.006. A_750/700_ denotes the maximum absorption ratio at *t* = 0 min. The dark reversion of AgP1 was similar to previously published results, where three components were used^34^. DrBphP however had a faster dark reversion rate and different time constants than reported before^35,71^. We therefore note that the differences in experimental parameters, including the buffer system, may cause the discrepancies in the reversion rates. Although the comparison between other studies is therefore problematic, the reversion rates within this study are fully comparable. Addition of the DrRR affected only the third time constant of the DrBphP reversion, from 480 minutes to 140 minutes and their relative amplitudes, which means that the structural properties of the DrBphP that are responsible for the third reversion component is affected by the DrRR binding. **B.** Size-exclusion chromatography of DrBphP and AgP1 and their response regulators as EGFP fusions. In order to resolve the phytochromes from response regulators, we fused the response regulators with EGFP. These EGFP-RR proteins could be exclusively detected with 509 nm emission, whereas phytochrome retention could be detected with biliverdin-specific 700 nm absorption. Interacting proteins from *D. radiodurans* (DrBphP and EGFP-DrRR) are in left, and the proteins from *Agrobacterium fabrum* (AgP1, EGFP-AtRR) are in right. BphP retentions are plotted at 700 nm absorption (upper panels) and EGFP-RR retentions are plotted as 509 nm fluorescence (lower panels). **C-D**. Size-exclusion chromatography of phytochromes and their response regulators in two different buffers: 30 mM Tris/HCl, pH 8.0 (C), and 67 mM phosphate, 200 mM NaCl, pH 8.0 (D). The experiments were conducted at 4°C with Superdex 200 Increase 3.2/300 (GE Healthcare). Protein retentions were detected at 280 nm absorption. The void volume (1.00 ml) was verified with Dextran, and the molecular weights in C were estimated from standard curve attained from Gel Filtration Calibration Kit LMW (GE Healthcare). Each protein was injected at 10 mg/ml concentration in 30 μl. The retention graphs show that DrRR remained monomeric in both conditions, whereas AtRR was mainly a dimer. DrBphP behaved like previously described^71^: Red light illumination led to a second phytochrome peak that elutes right after the main Pr peak (C). However, DrBphP readily oligomerized once NaCl is included, especially after red light illumination (D). Here, DrRR binding to DrBphP could not be detected in condition C. However, once DrRR is present in condition D, the DrBphP dimer peak is shifted and the oligomer peak disappeared, which points for DrBphP/DrRR interaction. In the case of AgP1, no changes in retention due to AtRR binding were detected. R = red light (655nm) illumination; FR = far-red light (785nm) illumination. Abbreviations: D = dark sample; R = red light (655 nm) illuminated sample.

**Supporting Figure S2.**
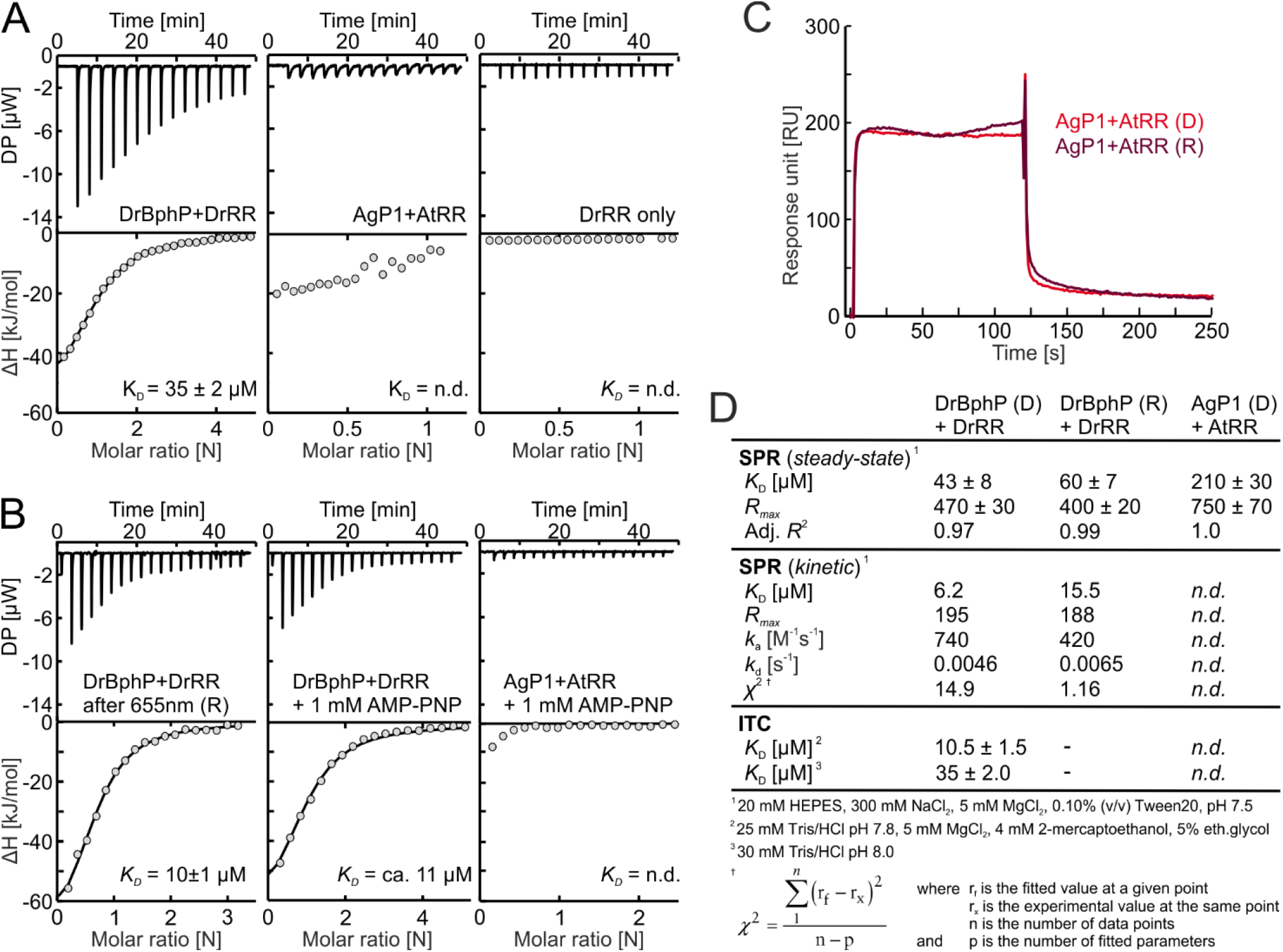
Additional measurements with isothermal calorimetry (ITC). **A**. DrRR binding to DrBphP measured in (30 mM Tris/HCl pH 8.0). DrBphP/DrRR interaction occurs with *K*_D_ of 35 ± 2 μM, and no AgP1/AtRR interaction was detected. **B**. DrBphP/DrRR interaction in buffer (50 mM Tris/HCl pH 7.8, 10 mM MgCl_2_, 8 mM 2-mercaptoethanol, 10% ethylene glycol) was unaffected by the red (655 nm) preillumination. This supports the result from SPR (Figure 2A) that illumination would not significantly affect the affinity. However, the ITC measurements have long pre-equilibration time (10 min) and slow acquisition time (~50 min), which leads to significantly reduced and constantly diminishing Pfr phytochrome population during the measurement. DrBphP/DrRR interaction and AgP1/AtRR interactions are unaffected by addition of 1 mM AMP-PNP in the buffer. **C**. Surface plasmon resonance (SPR) measurements of AgP1/AtRR interaction pair in dark (D) or after red light illumination (R). The measurement was conducted as in Figure 2A by applying AgP1 (100 μM) to AtRR-coupled sensor surface. **D**. Table summarizing the results from SPR and ITC analyses, along with additional fitting parameters. Adjacent R^2^-values indicate an agreement for the steadystate fits, with the best agreement approaching a value of 1.0. The χ^2^ values close to 10 indicate a good kinetic *K*_D_ estimation in the case on DrBphP/DrRR interaction. *R_max_* values give information on the maximal SPR response in saturating protein concentrations.

**Supporting Figure S3.**
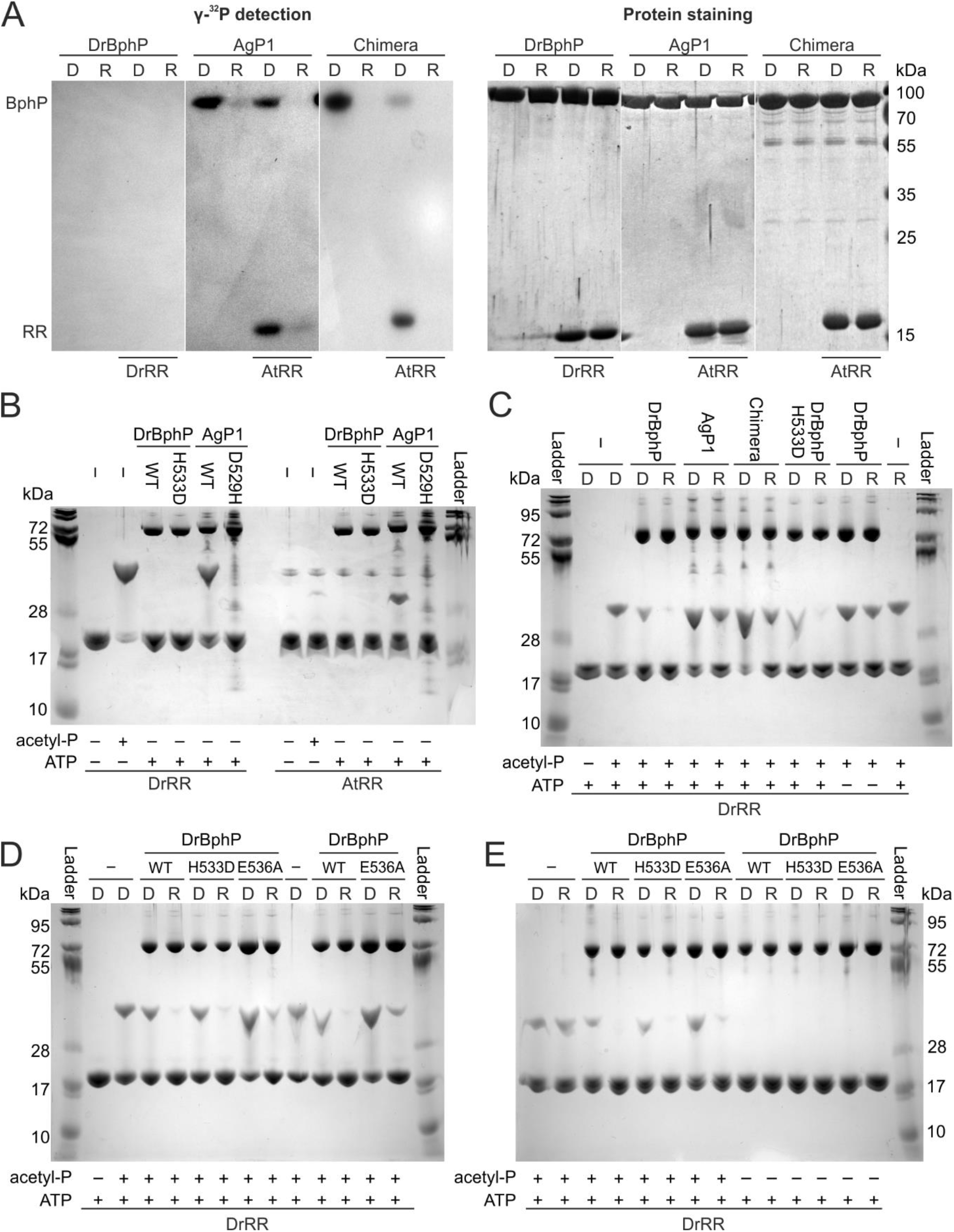
Full gels for radiolabeled and Phos-tag assays. **A**. Full gels from the radiolabeled ATP assay shown in the Figure 3A in the main text. **B**. Full gel of the kinase assay shown in Figure 3B. **C**. Full gel of the results shown in Figure 3C. The gel also shows that the phosphatase activity of DrBphP is stalled when ATP was excluded from the reaction. **D**. Full gel of the results shown in Figure 3D. **E**. Full gel of the results shown in Figure 3E. As seen in gels B and D, red light did not affect the stability of the free phospho-DrRR. The phytochrome samples are indicated above the gels. The response regulator, its treatment with acetyl phosphate treatment, and the inclusion of ATP are indicated below each gel. All measurements have been repeated at least three times, and not all repeats have been shown here.

**Supporting Figure S4.**
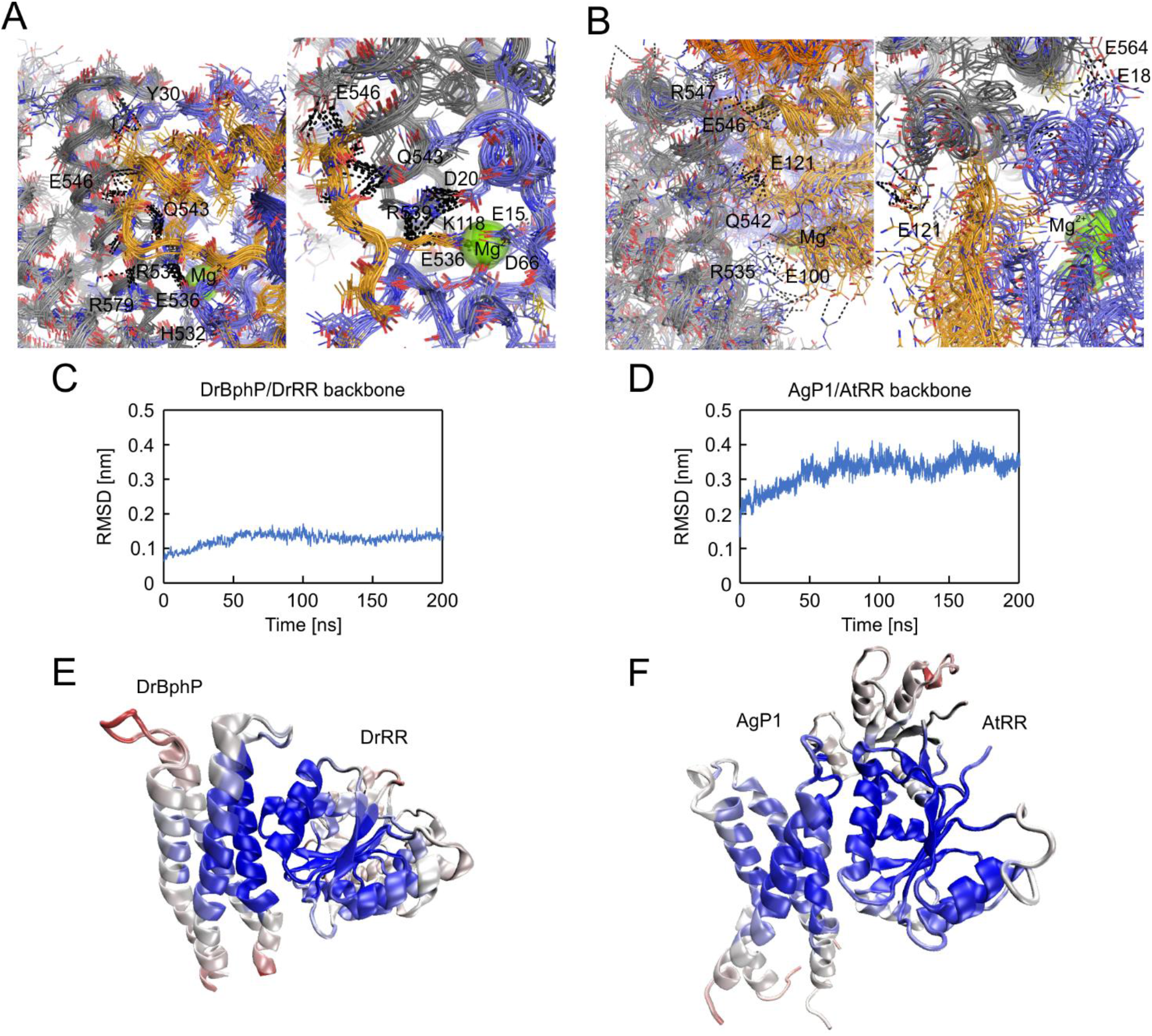
Flexibility of the HK/RR complexes in AgP1 and DrBphP. **A–B**. Relaxed solvated models of complexes DrBphP/DrRR (A) and AgP1/AtRR (B). 11 representative models are shown after global structural alignment by using PyMOL Molecular Graphics System version 2.3.3 (Schrödinger, LLC). Some of the polar interactions and salt bridges have been indicated. Molecules have been colored as in Figure 5 of the main text. **C–D**. RMSD of the backbone atoms along the 200 ns MD trajectory of DrBphP/DrRR complex (C) and AgP1/AtRR complex (D). **E–F**. Color maps of the amino acids heavy atoms root-mean square fluctuations (RMSF) for DrBphP/DrRR complex (E) and AgP1/AtRR complex (F). Blue color denotes rigid regions, red color denotes more flexible regions.

**Supporting Figure S5.**
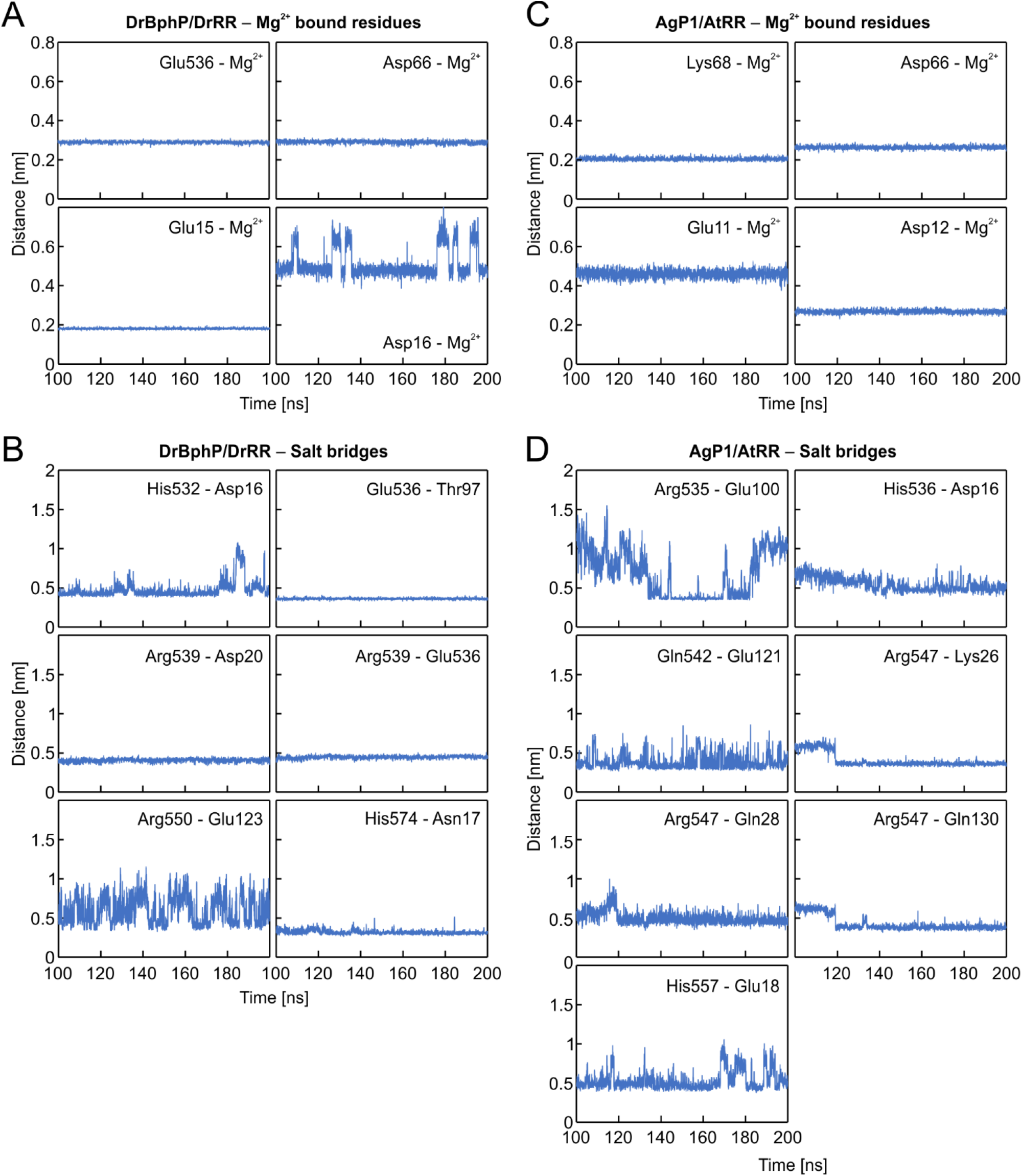
Distance plots of the structural parameters in HK/RR complexes. Each individual plot represents distance between the head groups of the residues at 100–200 ns in the MD trajectory after the initial 100 ns equilibration. **A**. Distances between Mg^2+^ ion and head groups of bound residues in DrBphP/DrRR complex. **B**. Salt bridges between DrRR and DrBphP proteins. **C**. distances between Mg^2+^ ion and head groups of bound residues in AgP1/AtRR complex. **D**. Salt bridges between AtRR and AgP1 proteins.

**Supporting Figure S6.**
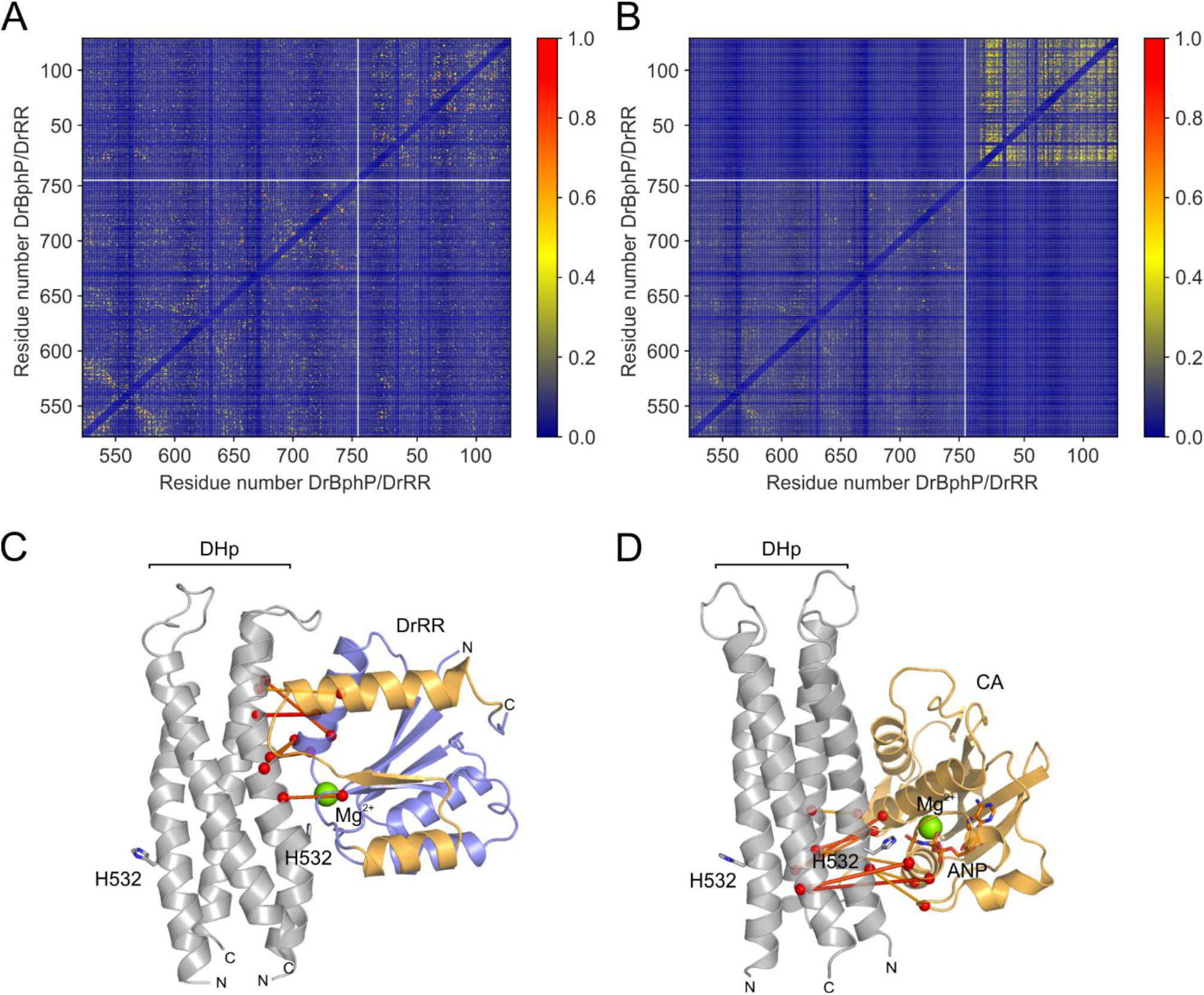
Covariance analysis of in the DHp/RR and DHp/CA interfaces. **A**. Covariance matrix of the tandem BphP-RR sequences. Blue color indicates low and red high pairwise covariance. **B**. Covariance matrix of scrambled sequences. Here, little covariance between the HK and RR sequence segments is visible, thus verifying the specificity of the results shown in A. **C–D**. Strong covariances between residues in the DHp helices and their cognate response regulators (RR), and a catalytic ATP-binding (CA) domains. The same models are shown as in Figure 5A and Figure 6A. The extent of the covariance is color-coded in such a way that the red lines indicate high covariance and yellow lines intermediate covariance. Only the covariance lines above a cutoff value of 0.6 are shown.

**Supporting Figure S7.**
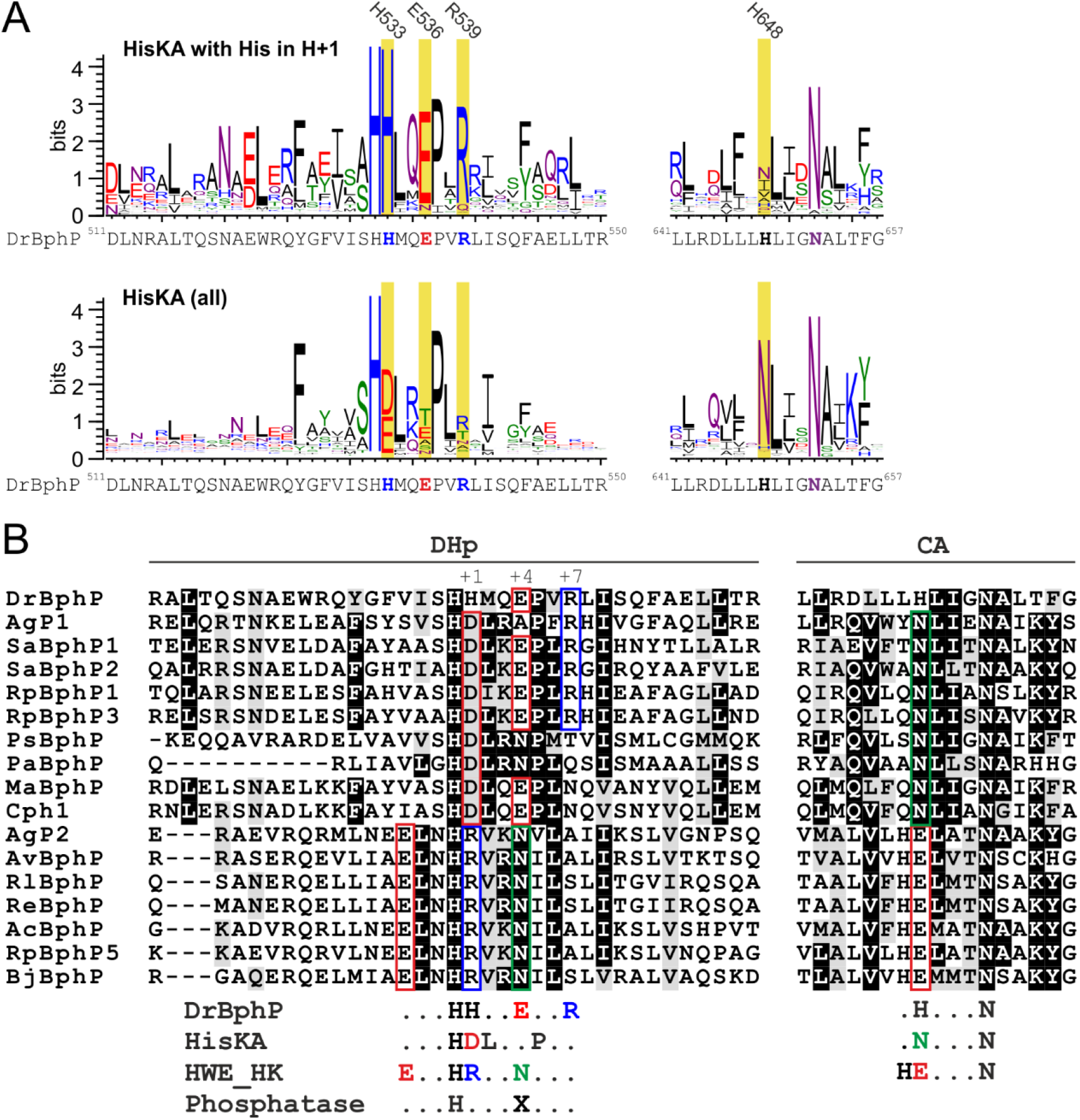
Sequence variation of the key catalytic residues in histidine kinases. **A**. Sequence logo showing the conservation of selected residues in the HisKA occurrences. Two occasions have been shown: One where the H+1 position in the DHp helix is occupied by a histidine (His533 in DrBphP), another where H+1 site is not restricted to as a histidine. The sequence logo in the former occasion were plotted from 192 sequences, which resulted noisier graph than in the latter occasion. In the case where H+1 is histidine, the conservation of the N-box Asn in the CA domain is lost (H648 in DrBphP). However, the occurrence of Glu in H+4 position as well as Arg in position H+7 is increased in these cases. The key residues are shaded in yellow. **B**. Sequence alignment of representative bacteriophytochromes. Below the alignment are shown the key DrBphP residues discussed in the main text, along with consensus sequence of HisKA, HWE_HK, and phosphatase. Alignment shows that the bacteriophytochromes above Cph1, and Cph1 itself, likely belong to HisKA family. Although DrBphP resembles other HisKA proteins, it misses a few functionally important residues. AgP1 has all the key residues required for HisKA activity. AgP2 and the phytochromes below it belong to HWE_HK proteins with a characteristic arginine residue in its H+1 position and a glutamate in the N-box of the CA domain. As for phosphatase activity, a threonine, asparagine or glutamate in H+4 position indicate that all selected phytochromes may also act as phosphatases. As a sole exception, the phosphatase activity of AgP1 may be hindered due to an alanine in this position. In conclusion, most phytochrome sequences have features that enable both kinase and phosphatase activity. Notable exceptions for this are AgP1 and DrBphP. Full-length sequences were aligned in Jalview 2.11.1.0^92^ usign ClustalO^93,94^ with standard settings. The amino acid conservation was visualized with Boxshade version 3.21. Uniprot accession numbers in the same order as in the alignment: DrBphP (*Deinococcus radiodurans* - Q9RZA4), AgP1 (*Agrobacterium fabrum* - Q7CY45), SaBphP1 (*Stigmatella aurantiaca* - Q097N3), SaBphP2 (*Stigmatella aurantiaca* - Q09E27), RpBphP1 (*Rhodopseudomonas palustris* - Q6N5G3), RpBphP3 (*Rhodopseudomonas palustris* - Q6N5G2), PsBphP (*Pseudomonas syringae* - Q885D3), PaBphP (*Pseudomonas aeruginosa* - Q9HWR3), MaBphP (*Microcystis aeruginosa* - B0JT05), Cph1 (*Synechocystis sp. PCC 6803* - Q55168), AgP2 (*Agrobacterium fabrum* - A9CI81), AvBphP (*Agrobacterium vitis* - B9K3G4), RlBphP (*Rhizobium leguminosarum* - Q1MCX7), ReBphP (*Rhizobium etli* - B3PX96), AcBphP (*Azorhizobium caulinodans* - A8HU76), RpBphP5 (*Rhodopseudomonas palustris* - Q6NB40), BjBphP (*Bradyrhizobium japonicum* - A0A023X9Y5).

## Notes

### Competing Interest Statement

The authors have declared no competing interest.

